# Comparative vitreous proteomic profiling of proliferative diabetic retinopathy and diabetic with no-retinopathy subjects implicates impaired autophagy in DR pathogenesis

**DOI:** 10.1101/2024.07.08.601678

**Authors:** Sarmeela Sharma, Shahna Shahul Hameed, Sushma Vishwakarma, Jay Chhablani, Mudit Tyagi, Raja Narayanan, Rajeev R Pappuru, Ghanshyam Swarup, Subhabrata Chakrabharti, Inderjeet Kaur

## Abstract

**Purpose:** Diabetic retinopathy (DR) is a neurovascular complication of diabetes (DM) causing the loss of neurons (ganglion cells) in the retina. The disease etiology and potential pathogenic mechanisms in this disease remains unclear. In the present study, we aimed to further understand the key and novel pathogenic mechanisms involved in DR pathogenesis by taking cues from our global proteomics data.

**Methodology:** The study was approved by the institutional review board (IRB) of LVPEI, Hyderabad, India. Vitreous humour samples (PDR; n=3, DM; n=3, Control; n=3) were collected from patients undergoing vitrectomy and subjected to LC-MS-MS analysis. The acquired raw data were searched against the human vitreous proteome and was further analysed by various bioinformatic and proteomic tools. Western blotting and IHC was performed to validate crucial pathways. Blood samples from patients (DM, PDR & NPDR) and controls (n=50); cadaveric retinas from diabetic and non-diabetic donors (n=10) and epiretinal membranes (ERM, n=10) from DR cases and controls were collected and RNA was isolated. Quantitative expression of genes involved in autophagy were performed. ɅɅCT was compared across different categories and significance estimated using a student t-test.

**Result:** A total of 1079 proteins were identified with 16 completely novel proteins in eye proteome. Top pathways identified were autophagy, inflammation, LXR/RXR activation (lipid metabolism), ROS generation by macrophages, apoptosis and protein degradation. Regulatory proteins identified were associated with cell death, phagocytic activation, angiogenesis and apoptosis. Autophagy inducers such as ROS was found to be accumulated in the DR vitreous. TREM2, microglial receptor was identified as a novel protein in PDR vitreous. The expression of *TREM2, an* autophagy-associated gene was significantly (p-value = 0.05) upregulated in all categories as compared to control (NDM and/or NDM/No-DR). TREM2 protein also seemed to colocalise with microglial marker F4/80 in retinal tissues and intense expression was observed near the blood vessels in case of PDR retina. Other autophagy-associated markers were also differentially regulated in DR as compared to controls.

**Conclusion:** This study emphasises on the strong role of autophagy pathways and its associated genes in the development of DR.

## Introduction

Diabetic retinopathy (DR) is one of the major sight-threatening retinal neurovascular complications of diabetes with a global prevalence of 34.6% and 21.7% in India (Gadkari et al., 2016). Even though the clinical features of this disease such as microaneurysm, edema, vitreous hemorrhage and retinal detachment are well known, the underlying biochemical changes occurring in the disease pathogenesis is least understood yet due to its extreme metabolic complexity (An D et al., 2022). The global proteome profiling can provide an overview of the key mechanism involved in such complex diseases by providing an association of altered proteins with disease pathogenesis, further an in-depth analysis of these could help to identify the potential therapeutic target. Vitreous humor is adhered to the retina and the protein shed into the vitreous from a normal physiological or pathological state would be retained until it is surgically removed, thus vitreous proteome studies provide a snapshot of the overall events happening in the retina (Monteiro JP et al., 2015). Proteome profiling using the mass-spectrometry based approach has been widely used for addressing key players and pathways involved in various retinal complexities by taking vitreous humor as a reliable source tissue (Dos Santos FM *et al* 2022).

Several groups have done global vitreous proteome profiling to catalogue the proteins under a normal physiological or with respect to a pathological state using multiple strategies (Aretz, Krohne et al., 2013, Murthy, Goel et al., 2014, Nobl, Reich et al., 2016, Santos, Gaspar et al., 2018, Zou, Zhao et al., 2018). Though these studies used similar source tissue for proteome profiling, a huge variability was noted in the identified proteins and in their numbers even between similar diseases phenotypes due to the variations in their study designs. For example, Wu *et al* in 2016 have profiled proteome retinal detachment cases using iTRAQ combined with nano-LC-ESI-MS/MS and found a total of 750 proteins from 16 vitreous samples (Wu, Ding et al., 2016), while, Santos *et al* in 2018, catalogued 1030 proteins from 8 –vitreous samples of retinal detachment cases using LC-ESI-MS/MS-iTRAQ (Santos et al., 2018). Likewise, multiple vitreous proteome studies were conducted to explore the underlying disease mechanism of DR. The first vitreous proteome analysis for DR was performed by Nakanishi *et al*. in 2002 that identified proteins such as complement C4, α1-antitrypsin, α2-HS glycoprotein and pigment epithelium-derived factor (PEDF) in PDR vitreous for their potential role in disease pathogenesis (Nakanishi, Koyama et al., 2002). This was followed by many other proteomics studies that explored the disease mechanisms in PDR and suggested the involvement of various proteins, such as apolipoproteins, complement proteins, zinc α-2 glycoproteins, serotransferrin, coagulation pathway proteins (Garcia-Ramirez, Canals et al., 2007, Wang, Feng et al., 2013), kallistatin, thioredoxin, von Willebrand factor (vWF), chromogranin (Kim, Kim et al., 2010, Kim, Kim et al., 2007), proteins of the complement and coagulation system, glutathione peroxidase 3, Immunoglobulins and cellular adhesion molecules (Hernandez, Garcia-Ramirez et al., 2013, Loukovaara, Nurkkala et al., 2015) in disease pathogenesis. Recent studies have highlighted involvement of extracellular remodelling, inflammation, neuroprotective pathway inhibition, complement, coagulation and perturbation in visual transduction pathway in DR (Li, Jin et al., 2021, Sen, Udaya et al., 2023, Weber, Zhao et al., 2021). This signifies the requirement of precise optimizations of the experimental designs at every stage i.e. from sample preparation to mass spec type to obtain a reliable and reproducible data with maximum number of protein identification, especially when dealing with complex metabolic diseases like DR.

While many studies have explored the disease mechanisms in DR by cataloguing the proteins, many proteins and their associated pathways remain unidentified. Studies have mainly focused on the angiogenic and inflammatory aspects of DR pathology. However, the interplay of neurodegeneration and neuroinflammation is yet to be elucidated. Therefore, our study focused on this aspect of DR pathology. Taking cues from proteomics data, we have further explored the role of neurodegeneration with focus on autophagy and neuroinflammation in DR pathogenesis.

## Materials and Methods

### Enrolment of study participants

The study participants were recruited from Smt. Kannuri Santhamma Centre for vitreoretinal Diseases, L V Prasad Eye Institute, Hyderabad, India. The study was done according to the guidelines of Declaration of Helsinki and study was approved by Institutional Review Board (LEC-BHR-P-04-21-626). Vitreous samples were collected from control and diabetic with no retinopathy subjects while undergoing surgeries for idiopathic macular hole (IMH).

### Sample preparation and Mass spectrometry

200-300 µl samples were collected in a cryovials at ice cold condition and centrifuged at 14,000rpm for 10 minutes to remove the cellular debris. The samples were lysed using equal volume of RIPA buffer followed by overnight precipitation with ice-cold acetone at -80° C. The samples were centrifuged at 14,000 rpm for 1 hour and solubilized the protein in 1X PBS and quantified using BCA assay .100µg of protein from controls and patients were loaded on to a 10% SDS PAGE followed by staining with Coomassie brilliant blue staining. After the de-staining in-gel trypsin digestion was performed and samples were loaded into Mass spectrometry using Q Exactive (Thermo Scientific) interfaced with nanoflow LC system (Easy nLC 1200, Thermo Scientific). Thermo Xcalibur MS data system software was used as the system controller. Trypsin-digested desalted peptides (10μL) were loaded onto PepMap RSLC C18 3 μm, 100 Å, 75 μm × 15 cm (Thermo Fisher Scientific) at a flow rate of 1 μL/min. Peptides were separated using 120 min linear gradient of the mobile phase [5% ACN containing 0.1% formic acid (buffer-A) and 95% ACN containing 0.1% formic acid (buffer-B)] at a flow rate of 300 nL/min. The data acquisition was performed in the electrospray ionization positive mode, with charge state + 2, capillary temperature 300 °C and spray voltage at 2.2 kV. Full scan MS spectra were acquired at a resolution 70,000 and scan range 400– 1750 m/z. The top 10 most intense peaks containing doubly, and higher charged states were selected for sequencing and fragmentation in the HCD (High Energy Collision-Induced Dissociation) mode at a scan resolution of 17500 with normalized collision energy set to 29%. Dynamic exclusion was activated to minimize repeated sequencing for the entire sequencing event.

### MS/MS data analysis

The acquired raw data for vitreous humour samples were searched against the human proteome from Uniprot database (release 2018.09 with 73099 entries) and a database of known contaminants using the Andromeda search engine and Maxquant (version 1.3.0.5). All MS/MS spectra from each sample were analysed using minimum one unique peptide for identification and 0.05% FDR (false discovery rate) on both peptide and protein level. Other search parameters included constant modification of cysteine by carbamidomethylation, enzyme specificity trypsin, Label-Free Quantification (LFQ) selected and match between runs with 2 min time window. iBAQ option was selected to compute abundance of the proteins.

### Bioinformatic analysis

The identified protein lists from the samples were further analysed by various bioinformatics tools for proteome analysis. The proteins were compared with eye proteome database for identifying the novel vitreous eye proteins detected from the present study. Abundant vitreous proteins were evaluated in the vitreous samples by analysing the top 20 proteins based on their intensity and peptide detected. Gene Ontology (GO) was performed for the hierarchical classification of identified proteins based on their involvement in major GO-Slim biological process, GO-slim molecular function and GO-Slim cellular components using PANTHER-GO database for the complete set of proteins in each category. Label free quantitation was done to evaluate protein expression based on their intensity. Differential expressions of proteins in diabetes compared to non-diabetic group and retinopathy compared to no-retinopathy groups were evaluated by calculating the mean the fold change.

### Network and Pathway analysis

Qiagen Ingenuity Pathway Analysis (IPA) software was used to find possible networks and pathways based on the differentially expressed proteins.

### Enzyme linked immunosorbent assay (ELISA)

ELISA was performed to assess the expression of LRG1 in an increased cohort (n=10 each category of PDR, DM and controls). Inflammation, autophagy and angiogenic pathways were validated by validation of their major pathway proteins (MMP-9, IL-8, VEGF, VEGFR1, LIPOCALIN-2, THBS2, PDGF-AA and Endothlein-1) using assays Bibiotech technologies India Pvt. Ltd. ELISA was performed in 25 samples each from PDR and controls. Vitreous protein samples were evaluated using the Luminex xMAP Immunoassay Kit (Thermo Fisher Scientific, USA), performed according to the manufacturer’s instructions. Vitreous samples were diluted (1:5) in the diluent provided in the kit. 50 μL of standards and samples were added to 96 well plates followed by the addition of 50 μL of diluted microparticle cocktails. Plates were incubated for 2 h at room temperature on a shaker at 800 rpm. Thereafter, the plate was washed thrice with freshly prepared 100 μL of 1× wash buffer. Next, 50 μL of biotinylated antibody cocktail was added to each well, following which the plates were sealed incubated for 1 h on a shaker at 800 rpm. After washing thrice, 50 μL of phycoerythrin-conjugated streptavidin was added to each well and the plates were again incubated at room temperature for 30 min on shaker at 800 rpm. After subsequent washing, 100 μL of wash buffer was added to each well and incubated for 2 minutes at room temperature on shaker at 800 rpm followed by measurement of fluorescence using a Luminex 100/200 system. Sample concentrations for each analyte were obtained based on the median fluorescence intensity relative to standards.

### Western blotting

Western blotting was performed for collagen-II and myocilin expression in the vitreous samples (n=10 each category of PDR and controls). Briefly, the processed vitreous samples were precipitated using acetone method of protein precipitation. Proteins were quantified using Micro BCA^TM^ protein assay kit (catalog no. 23235). 20 and 30 ug of protein was loaded onto 8 and 12% SDS-PAGE gel respectively for ColIIa and Myocilin protein quantification. After sufficient run, the proteins were transferred onto PVDF membrane. The membranes were blocked with Odyssey blocking solution from LICOR biosciences (928-40000). Membranes were incubated with primary antibodies Anti-mouse ColIIa (1:500, ab34712) and Anti-Rabbit Myoc (1:200, overnight at 4°C, ab41552. The next day, both the membranes were incubated with subsequent fluorescent labelled secondary antibodies at room temperature for 1 hour. This was followed by washing with 1xPBST and 1xPBS thrice and then visualized Odyssey Infrared Imaging System from Licor biosciences.

### Trichome Staining

Trichome Staining of retinal tissue: Sections were deparaffinized and hydrated using distilled water. Bouin’s fixative solution was used for fixing overnight at room temperature. Next day sections were washed in running water followed by a washing step with distilled water. Further sections were stained with Weigert’s iron hematoxylin solution for 10 minutes at room temperature. Again, washing was done under running water followed by a distilled water wash.

Sections were next stained with Biebrich scarlet acid fuchsin solution for 15 minutes. After distilled water wash, sections were differentiated in phosphomolybdic-phosphotungstic acid solution for 15 minutes. Next aniline blue solution was used for counterstaining for 1 minute. Lastly, sections were dehydrated and cleared using 95 percent ethyl alcohol, absolute ethyl alcohol and xylene, twice each for 2 minutes each. Mounting was done using DPX and visualization was performed.

### Immunohistochemistry

Immunohistochemistry was performed to check for the localization of TREM2 protein in the retinal layers from NPDR, PDR and controls (n=1). Retinal tissues were collected from the patients undergoing routine evisceration surgeries in the operating rooms with an informed consent. Parafilm embedded sections were made from the collected tissues. The sections were de-paraffinized using 1X sodium citrate buffer. After antigen retrieval, sections were fixed with paraformaldehyde, and incubated with triton-X 100 (0.5%) for permeabilization. Following this, sections were blocked with 2% BSA. Sections were incubated with primary antibody TREM2 (catalog no. PA5-87933) in 1:100 dilution overnight at 4°C. The next day, both the sections were incubated with subsequent fluorescent labelled secondary antibodies at room temperature for 1 hour. This was followed by washing with 1xPBS thrice and then visualized under EVOS fluorescent microscope.

### Measurement of oxidative stress using DCFDA assay

To measure ROS stress, DCFDA assay was performed in 10 samples each from PDR and controls. 5uL of undiluted vitreous samples was added to a 96 well plate. 195uL of DCFDA dye (10 μM) prepared in 1xPBS was added to the samples and incubated at room temperature for half an hour. After half an hour, absorbance was measured using spectra max at excitation/emission spectra of 485/535nm of wavelength. The results were calculated, and the data was measured for statistical significance using the student’s t-test.

### Semi-Quantitative PCR

RNA from blood samples of DR patients was extracted by the organic method using TRIZOL reagent. Quality and quantity were assessed by agarose gel and nanodrop spectrophotometer respectively. 500 ng of RNA was further reverse transcribed to cDNA using the iScript cDNA synthesis kit (Bio-Rad, CA, USA). Semi-quantitative PCR was carried out using specific primers for the following genes; *TREM2, LC3-II, LDLR, APOE, ATG4b, ADAM10, CATSB, APOE, ATG4b, CASP8, P62* and *β-3TUB* while *β-actin* was used as an endogenous control.

### Establishing primary cell culture from retinal tissues

Primary mixed retinal cell (PMRC) culture was established from retinal tissue chunks collected from patients undergoing evisceration surgery. Retinal tissue was chopped and disaggregated enzymatically by trypsin digestion into cell suspension followed by removal of necrotic cells and fat. Cells were resuspended and grown in lysine coated plates containing fresh DMEM medium with 10% FBS and 1% Penstrep. After sufficient growth and sub-culture, at passage 2, the PMRCs established above was treated with D-Glucose 20 mM for 24 hours to mimic the hyperglycaemic stress.

### Gene expression analysis

After 24 hours, cells were collected, RNA was isolated and converted to cDNA. Quantitative Real-Time PCR was performed for following genes: *TREM2, LC3-II, LDLR, APOE, ATG4b, ADAM10, CATSB, APOE, ATG4b, CASP8, P62* and *β-3TUB* using SYBR green chemistry. Cells not treated with D-glucose (20mM) served as controls for the experiment.

#### Immunofluorescence assay

After 24 hours of high glucose treatment, cells were fixed with 4% formaldehyde followed by 0.5% triton x-100 permeabilization and blocking with 2% BSA for an hour at room temperature. Cells were then incubated with primary antibodies TREM2 (1:100) and LC3b (1:200) overnight at 4⁰C overnight. Afterward, Alexa Fluor–coupled secondary antibodies (1:300) raised in suitable host were added for an hour at room temperature. Cell nuclei were visualized with DAPI (1:200; Molecular Probes, Thermo Fisher Scientific USA). All high-resolution fluorescent images were captured at 20x using EVOS fluorescent microscope.

#### Statistical analysis

The gene expression data was compared among the different categories of DR compared to controls. The fold change was calculated, and significance was checked using the student *t-test*.

## Results

### Demographic of the study subjects

Demographic details for the study subjects used for global vitreous proteome analysis is given in table1. Vitreous humor samples were collected from patients with the proliferative stage of diabetic retinopathy (PDR: n=3), diabetic patients who did not develop any clinical features of retinopathy (NDR: n=3) and control from non-diabetic subjects without any retinopathy (NDM: n=3). All the samples were taken from same gender (male) and similar ethnicity. There was no significant-difference in age between the groups and none of the patients had received any intraocular anti-VEGF injections as a part of their ocular complications prior to the sample collection.

### Identification of proteins in vitreous samples by LC-MS/MS

Total concentrations of the proteins in the vitreous humor samples were calculated using BCA assay and almost similar mean concentration of proteins in each group were observed that ranged from 9.46±1.45 μg/μL in NDM, 8.83±3.97 μg/μL in NDR and 9.77±2.51 μg/μL in PDR. 100 μg of proteins from each of the samples were used for LC-MS/MS analysis without any abundant protein depletion method using an in-gel digestion method. This identified a total of 1079 proteins in all the four groups with FDR of 0.5% and minimum peptide for protein identification as two. A total of 959,773 and 760 proteins were identified in NDM, NDR and PDR respectively (Figure 1a). The qualitative comparative analysis of proteins was done in all the three groups using VENNY 2.1. This demonstrated 618 common proteins in all the four groups. 96 proteins were found to be commonly present only in NDM, NDR and PDR group. There were 175, 90 and 20 unique proteins that were found to be present in NDM, NDR and PDR respectively (Figure 1b). 54 proteins were found to be present only in NDM and NDR, 111 proteins were found to be commonly present only in NDM and PDR and 10 proteins were found to be commonly present only in the NDR and PDR Figure 1b).

**Figure 1.**
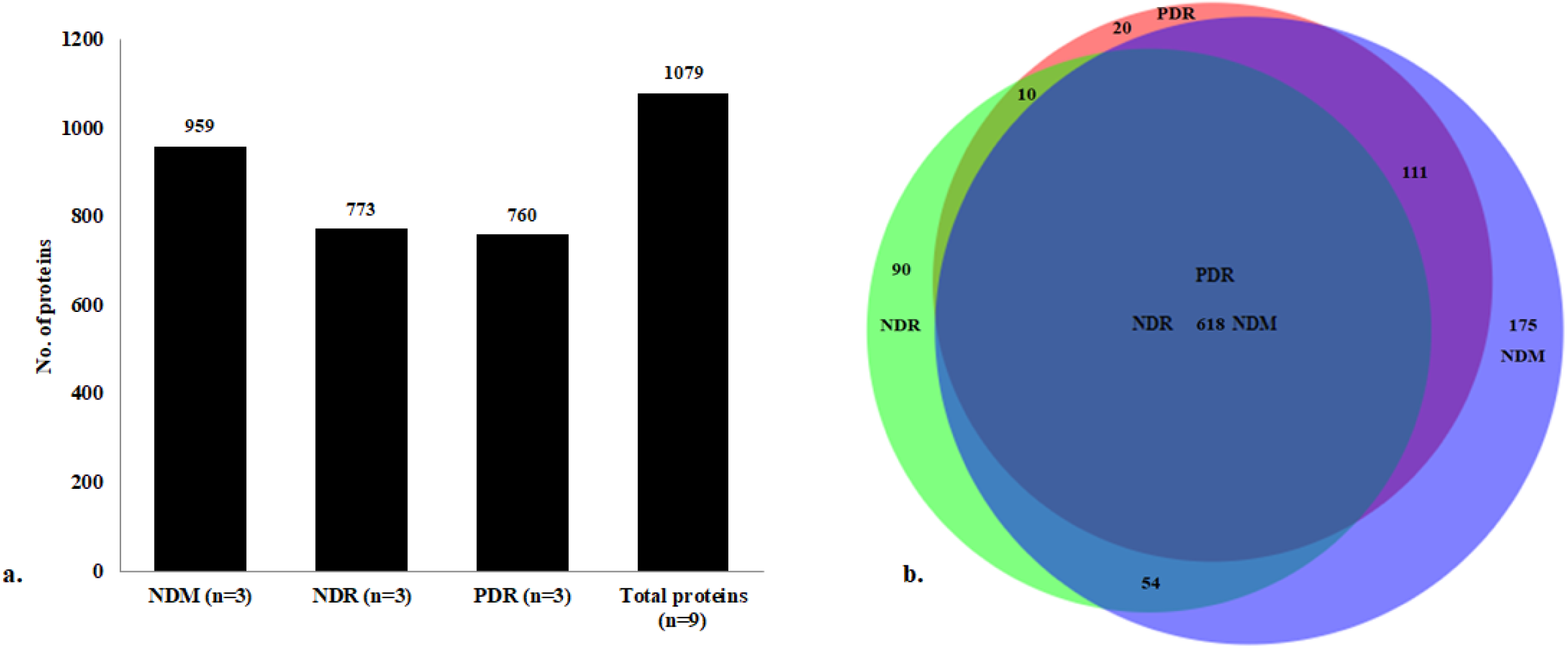
(a) Number of proteins present in each of the groups analyzed and the cumulative number of proteins in all the groups (PDR, NDR and NDM; n=3 in each group). (b) Venn diagram showing the overlap between proteins identified and unique proteins in the four different groups.

### Identification of the novel vitreous proteins

To identify the novel vitreous proteins from our dataset, we have compared the protein list from our data set with published vitreous protein database. It was observed that 938 proteins identified in the present study could be linked to the existing literature and databases, while 141 proteins were unique. But to verify these as novel proteins, we further implemented criteria such as presence of these proteins in any of the three independent vitreous samples with a minimum of 2 peptides and sequence coverage ≥2%. Further proteins with unmapped IDs were removed and this led to the identification of 27 novel vitreous proteins. Eleven of these proteins were found to be reported in eye proteome database (RPE-choroid, ciliary body, optic nerve, tear, sclera, iris, cornea, lens) and 16 proteins were not reported in any of the eye proteome database (Table 2 and 3).

**Table 1:** Demographics of study subjects used for global proteome profiling.

| Categories of patients | Age in Yrs. (Mean ±SEM) | Gender | DM Type | No. of DM Yrs. | Indication for surgery |
| --- | --- | --- | --- | --- | --- |
| NDM (n=3) | 58 ± 3.9 | M | N/A | N/A | Retinal Detachment |
| NDR (n=3) | 63 ± 5.1 | M | Type 2 | 4 ± 1.5 | Idiopathic macular hole |
| PDR (n=3) | 62 ± 4.1 | M | Type 2 | 19 ± 1 | Tractional Retinal Detachment |

**Table 2:** List of proteins identified in eye proteome database and their corresponding peptides and sequence coverage in the present cohort.

| Protein name and Gene ID | Mol. Wt. (kDa) | Seq. Length | NDM (Peptides, n=3) | NDM Seq. coverage (%) | NDR (Peptides, n=3) | NDR Seq. coverage (%) | PDR (peptides, n=3) | PDR Seq. coverage (%) | Tissue expression |
| --- | --- | --- | --- | --- | --- | --- | --- | --- | --- |
| Ig kappa chain V-I region HK101 | 12.73 | 117 | 2 ± 0.5 | 13.7 | 2 ± 0.5 | 13.7 | 1.3 ± 0.33 | 13.7 | Tear |
| Ig lambda chain V-VI region AR | 12.5 | 117 | X | X | 1.6 ± 0.3 | 29.6 | X | X | Cornea |
| Ig heavy chain V-III region KOL | 13.07 | 117 | 3.6 ± 0.3 | 34.2 | X | X | 2.3 ± 0.88 | 33 | Tear |
| Ig heavy chain V-III region JON | 12.9 | 117 | 16.7 ± 0.8 | 38.5 | 15 ± 2.6 | 35.3 | 15 ± 2.64 | 38.5 | Tear |
| Stromelysin-1 (MMP3) | 53.9 | 477 | 5 ± 3.2 | 6 | 2.6 ± 0.3 | 2.46 | X | X | RPE -choroid |
| Vesicle-associated membrane protein 1 (VAMP1) | 12.9 | 118 | 1.7 ± 0.8 | 9 | 5.3 ± 0.6 | 13.6 | 3 ± 1.52 | 13.6 | Retina, cornea, Iris, Ciliary body, optic nerve |
| HLA class I histocompatibility antigen, Cw-12 alpha chain (HLA-C) | 40.8 | 366 | 5 ± 1.1 | 9.9 | 6.6 ± 0.3 | 18.8 | X | X | Retina, cornea |
| Histone H2B type F-S (H2BFS) | 13.9 | 126 | 10 ± 5.5 | 32 | 7 ± 1.1 | 23.3 | 5 ± 2.08 | 17.7 | Retina, RPE-choroid |
| Histone H3.1 (HIST1H3A) | 15.4 | 136 | 6.6 ± 4.2 | 16.2 | 4.3 ± 1.2 | 16.4 | 3.6 ± 1.66 | 13 | Retina, cornea, Ciliary body, RPE-Choroid, lens |
| Protein FAM180A (FAM180A) | 19.7 | 173 | 7 ± 1.15 | 4.6 | X | X | X | X | Sclera |
| HLA class I histocompatibility antigen, Cw-17 alpha chain (HLA-C) | 41.2 | 372 | 4.6 ± 0.33 | 8.5 | X | X | 6.6 ± 1.2 | 4.6 | Retina |

**Table 3:** Novel vitreous proteins detected in the present study, the number of peptides identified and sequence coverage.

| Protein name and Gene ID | Mol. Wt. (kDa) | Seq. length | NDM (Peptides, n=3) | NDM Seq. coverage (%) | NDR (Peptides, n=3) | NDR Seq. coverage (%) | PDR (peptides, n=3) | PDR Seq. coverage (%) |
| --- | --- | --- | --- | --- | --- | --- | --- | --- |
| Ig heavy chain V-I region EU | 12.659 | 117 | 4.3 ± 1.3 | 25.3 | 3.3 ± 0.33 | 22.2 | 4.33 ± 1.51 | 25.3 |
| Ig lambda chain V-II region MGC | 12.382 | 118 | 12 ± 2 | 16.4 | 13 ± 1.73 | 17.8 | 13 ± 2.1 | 16.4 |
| Ig lambda chain V-II region TOG | 12.597 | 120 | 4 ± 1.52 | 13.3 | 7.6 ± 2.72 | 13.3 | 4.33 ± 1.8 | 13.3 |
| SEC14 like lipid binding 1 (SEC14L1) | 16.721 | 143 | 11.3 ± 1.2 | 7.7 | 13 ± 1.73 | 7.7 | 16.33 ± 1.4 | 7.7 |
| Putative neutrophil cytosol factor 1B (NCF1B) | 44.816 | 391 | 3 ± 1.5 | 4.26 | 4 ± 0.5 | 4.8 | 3 ± 1.62 | 4.26 |
| GTPase IMAP family member 4 (GIMAP4) | 39.033 | 343 | 2.6 ± 1.66 | 7.86 | 1.3 ± 0.33 | 7.7 | 3 ± 2 | 8.5 |
| Triggering receptor expressed on myeloid cells 2 (TREM2) | 25.447 | 230 | 1.3 ± 0.33 | 7.96 | 2 ± 0.57 | 10.2 | 3 ± 0.57 | 12.2 |
| Sulfotransferase family cytosolic 2B member 1 (SLUT2B1) | 41.307 | 365 | 3 ± 0.57 | 5.2 | X | X | 4.6 ± 0.33 | 5.2 |
| Mediator of RNA polymerase II transcription subunit 30 (MED30) | 20.277 | 178 | 8.6 ± 2.3 | 5.1 | 3.6 ± 0.33 | 5.1 | 6.3 ± 1.2 | 5.1 |
| N-acetyl neuraminate lyase (NPL) | 35.162 | 320 | 3.3 ± 2.02 | 8.96 | 2.6 ± 0.33 | 5.6 | 1.6 ± 0.33 | 5.6 |
| Zinc finger protein 268 (ZNF268) | 108.37 | 947 | 3.6 ± 0.33 | 2.4 | X | X | 3.6 ± 0.66 | 2.9 |
| Coiled-coil domain-containing protein lobo homolog (CCDC135) | 103.5 | 874 | 6.6 ± 0.88 | 4 | X | X | 6 ± 1.5 | 3.3 |
| 40S ribosomal protein S15a (RSP15A) | 27.168 | 237 | 2.3 ± 1.33 | 8.43 | 1 ± 0.57 | 2.3 | 2.3 ± 1.3 | 8.4 |
| Eukaryotic translation initiation factor 5A-1 (EIF5A) | 20.503 | 186 | 2.3 ± 0.66 | 10.763 | X | X | 2.6 ± 0.88 | 15.4 |
| Oxysterol-binding protein-related protein 6 (OSBPL6) | 106.3 | 934 | $2.3 \pm 0.33$ | 2.13 | X | X | $2 \pm 0.57$ | 1.5 |
| immunoglobulin lambda constant 2 (IGLC2) | 11.293 | 106 | $63.6 \pm 2.3$ | 89.93 | $63 \pm 7.2$ | 89.9 | $63.3 \pm 2.47$ | 89.9 |

### Enrichment of the proteins by Gene Ontology (GO)-slim analysis

Further to depict the overall understanding of the proteins present in each of the group, we have done GO-slim classification under the categories such as biological process, molecular function and cellular components using PANTHER (PANTHER, version 14.1) with Bonferroni correction and binomial testing. The PANTHER mapped 831, 677, 657 proteins from NDM, NDR and PDR respectively. Based on the mapped IDs and corresponding number of proteins in each subcategory percentage was calculated. For an optimal comparison, the significant biological process, molecular function and cellular compartments that involved >5% of the proteins in each of these three groups were analyzed.

### GO-Slim Biological process (GO-slim BP)

Overall, 34 significant GO-slim biological processes were identified in the NDM, NDR and PDR groups. Almost 40% of the proteins fell in the unclassified category under in each group (NDM: 307 (37%), NDR: 262 (38.8%) and PDR: 242 (36.9%)). As shown in figure 2. a large percentage of proteins were found to align with the biological process such as metabolic process (NDM: 33.7 %, NDR: 31.3%, PDR: 32.5%), signal transduction (NDM: 14.7 %, NDR: 13.9%, PDR:15.8%), cellular component organization or biogenesis (NDM: 16.04 %, NDR: 14.39%, PDR: 13.8%), immune system process (NDM: 10.37 %, NDR: 11.27%, PDR: 12.67%) and immune response (NDM: 8.56 %, NDR: 9.79%, PDR: 10.68%).

**Figure 2a.**
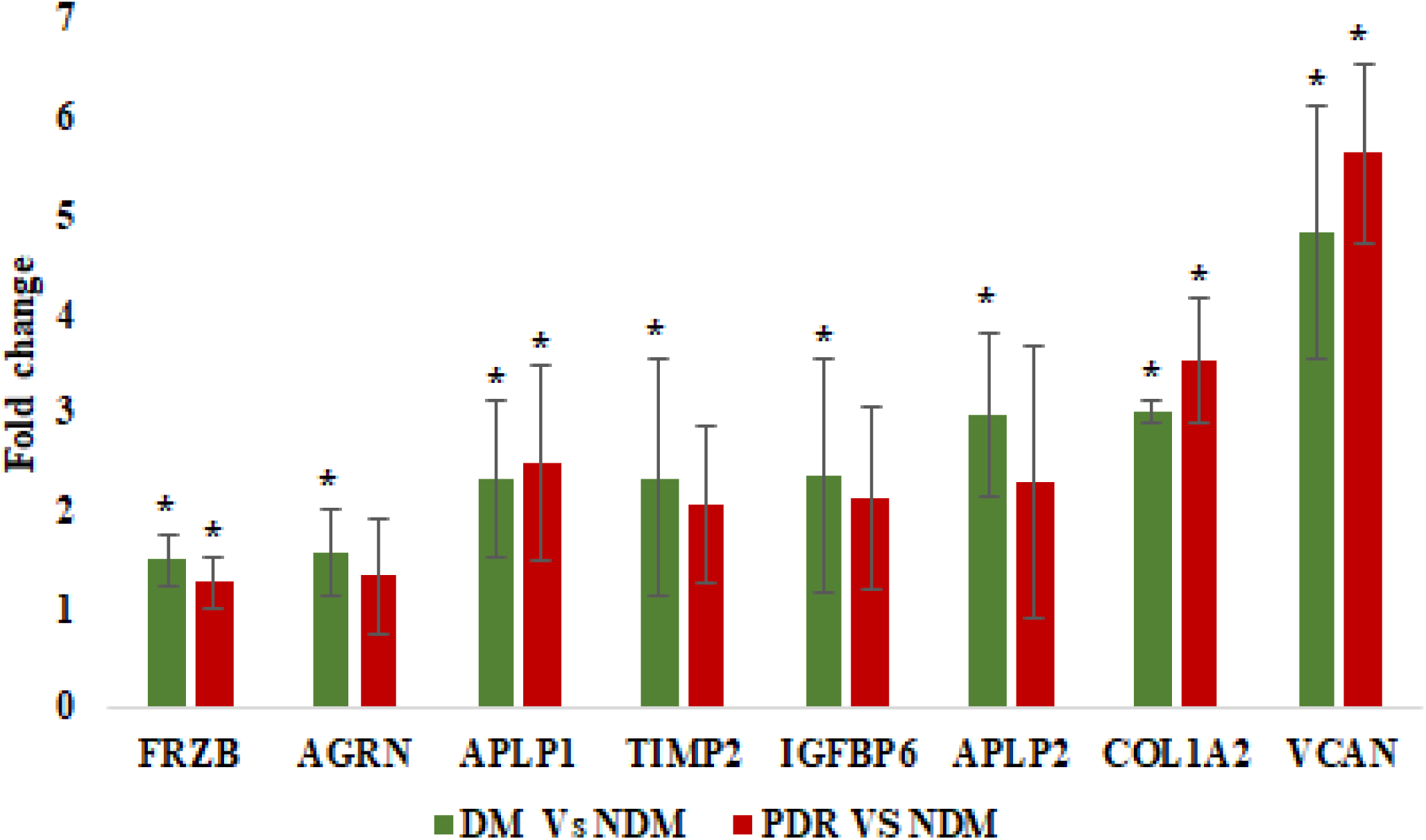
Significantly upregulated proteins in DMs vs NDM and PDR vs NDM group, \**p*<0.05

**Figure 2b.**
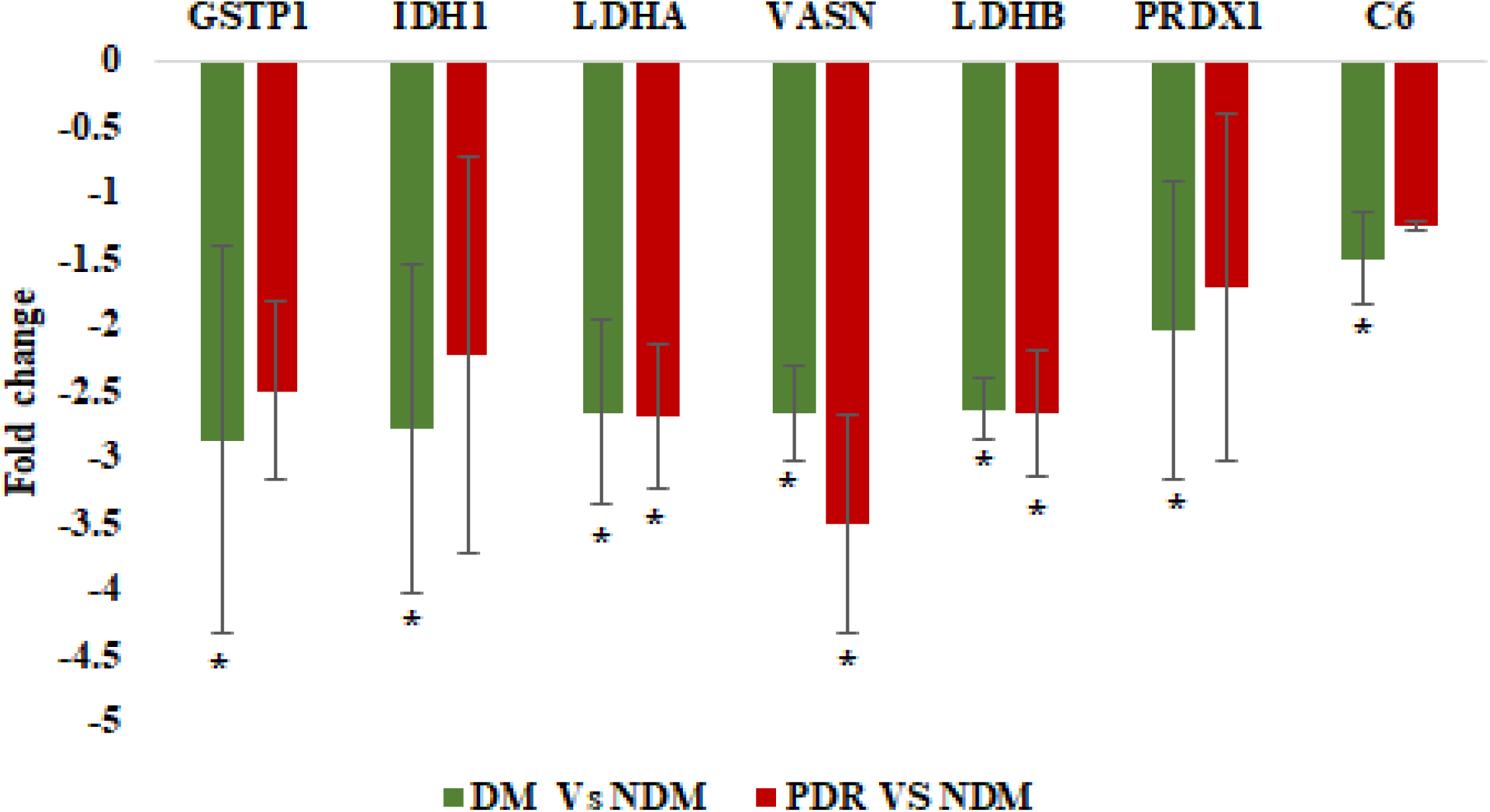
b. Significantly down-regulated proteins in DM vs NDM and PDR vs NDM group, \**p*<0.05

### GO-Slim Molecular function (GO-slim MF)

GO-slim molecular function identified 30 molecular functions in the NDM group, 28 molecular functions in NDR group and 29 in PDR groups. Like biological process, many proteins fell in the unclassified category (NDM: 380 (45.8%), NDR: 323 (47.92%), PDR: 315 (48.09%). The top 5 molecular functions identified were binding (NDM: 28.7%, NDR: 28.9%, PDR: 28.5%), catalytic activity (NDM: 28.46%, NDR: 27.15% PDR: 27.17%), protein binding (NDM: 20%, NDR: 21.9%, PDR: 21.06%), hydrolase activity (NDM: 15.9%, NDR: 16.02%, PDR: 16.48%) and receptor binding (NDM: 8.8%, NDR: 10.9%, PDR: 10.38 %).

### GO-slim Cellular component (GO-slim CC)

A total of 20 different cellular components were identified in proteins of PDR and NDM, while 19 were identified in the NDR group. A large proportion of the proteins were extracellular proteins that included proteins in the extracellular region (NDM: 20.98%, NDR: 26.40% PDR: 27.06%), extracellular region part (NDM: 19.54%, NDR: 24.4% PDR: 25.22%) and extracellular space (NDM: 17.73%, NDR: 22.25% PDR: 22.62%). The percentages of extracellular proteins were found to be higher in the PDR and NDR group compared to the controls. The proteins of the cytoplasmic part were found to be higher in NDM (10.61%) compared to NDR (6.37%) and PDR (7.95%).

Analysis of significantly expressed proteins in retinopathy progression independent of diabetes.

The understanding of differentially regulated proteins in each group is essential for disease characterization and for the identification of potential disease specific proteins. To achieve this, a proper understanding of disease induced alteration in the vitreous proteome is necessary. Hence, the mean fold change of proteins expressions based on the normalized intensity values were calculated in DM (PDR+ NDR) compared to NDM and retinopathy (PDR) compared to no-retinopathy (NDR+NDM). This approach helped us to identify protein alteration exclusively occurring in diabetes and in proliferative stage of the disease. Further, PDR to NDR and PDR to NDM were also calculated and compared with the values of diabetes and retinopathy induced changes. Proteins were considered as significant if their log2 fold intensity value were shown a minimum of 1.5-fold change with a *p*-value ≤ 0.05. The lists of the proteins identified in each of the categories are given in the Figure 2a. Also, proteins showing huge standard deviation were also removed to avoid any type of disparity. There were15 proteins that were found be differentially expressed in the DM group compared to NDM with a minimum fold change of 1.5 (*p*-value <0.05). These include 8 upregulated and 7 downregulated proteins. The levels of these proteins in other combinations were calculated to understand if the expressions of these proteins were further altered at proliferative stages of the disease. The down regulated proteins were majorly the proteins involved in detoxification mechanism and glycolytic pathway proteins. The expressions of these proteins were similarly maintained in the DM vs no-DM and PDR vs NDM, which proves diabetes is the key factor involved in these proteins downregulation and is not further affected by the proliferative changes occurring in the disease stage (Figure 2b).

Likewise, 4 upregulated and 6 downregulated proteins were found to be differentially regulated in the retinopathy stage. Level of these proteins’ expression were also evaluated in PDR vs NDR, which further clearly identified a similar protein expression as seen retinopathy vs no-retinopathy group. This further confirmed the expression of these 10 proteins were typically dependent on the advanced stage of the disease. Most importantly, VASN expression was also found to be downregulated significantly in DM compared to non-DM as well as PDR Vs NDM group. Also, a significant upregulation in the angiogenic protein LRG1 and downregulation of detoxification protein SOD3 was observed at retinopathy group along with other proteins (Figure 3a. and 3b).

**Figure 3a.**
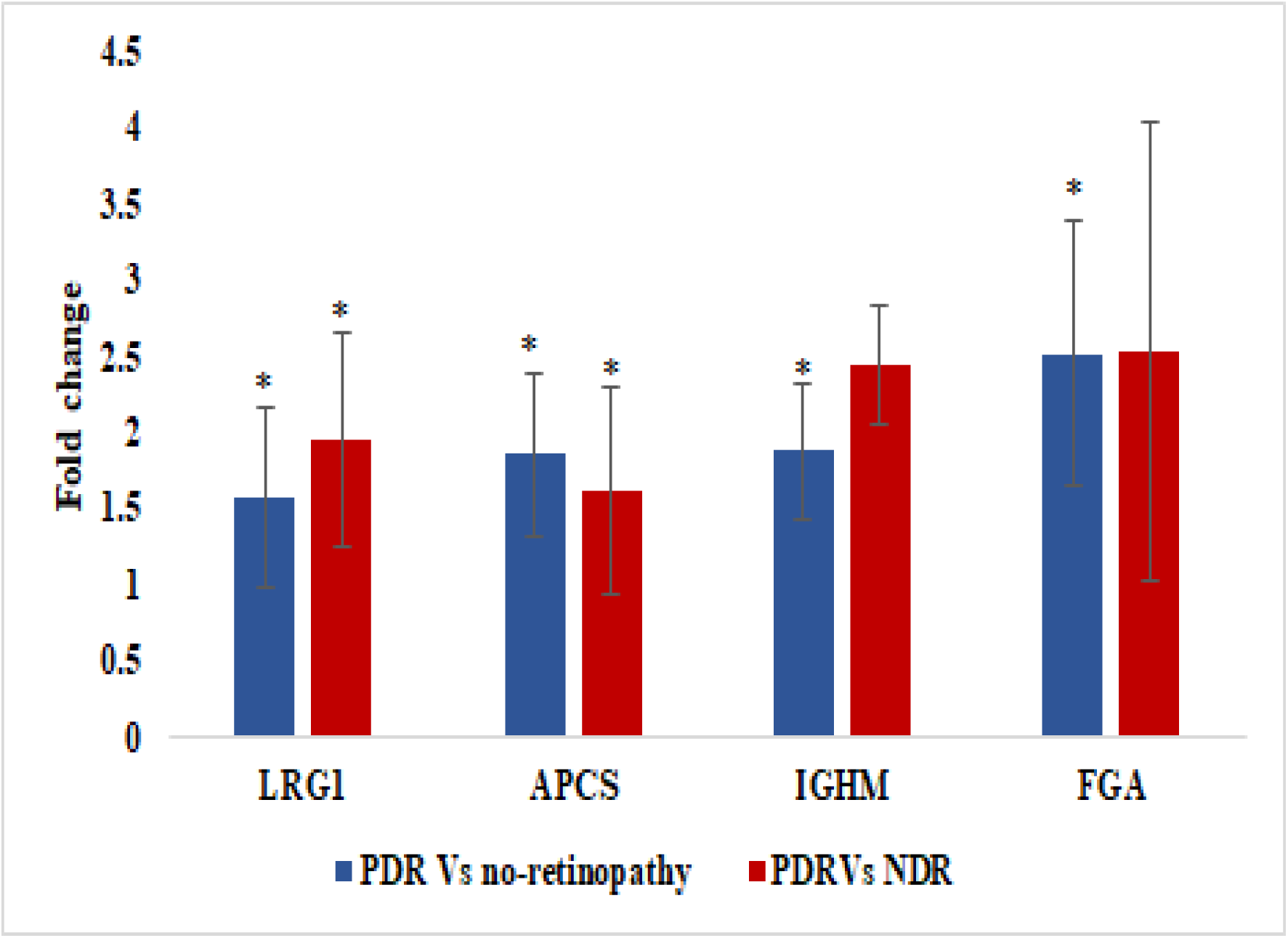
Significantly up-regulated proteins in PDRD vs no retinopathy groups, \**p*<0.05

**Figure 3b.**
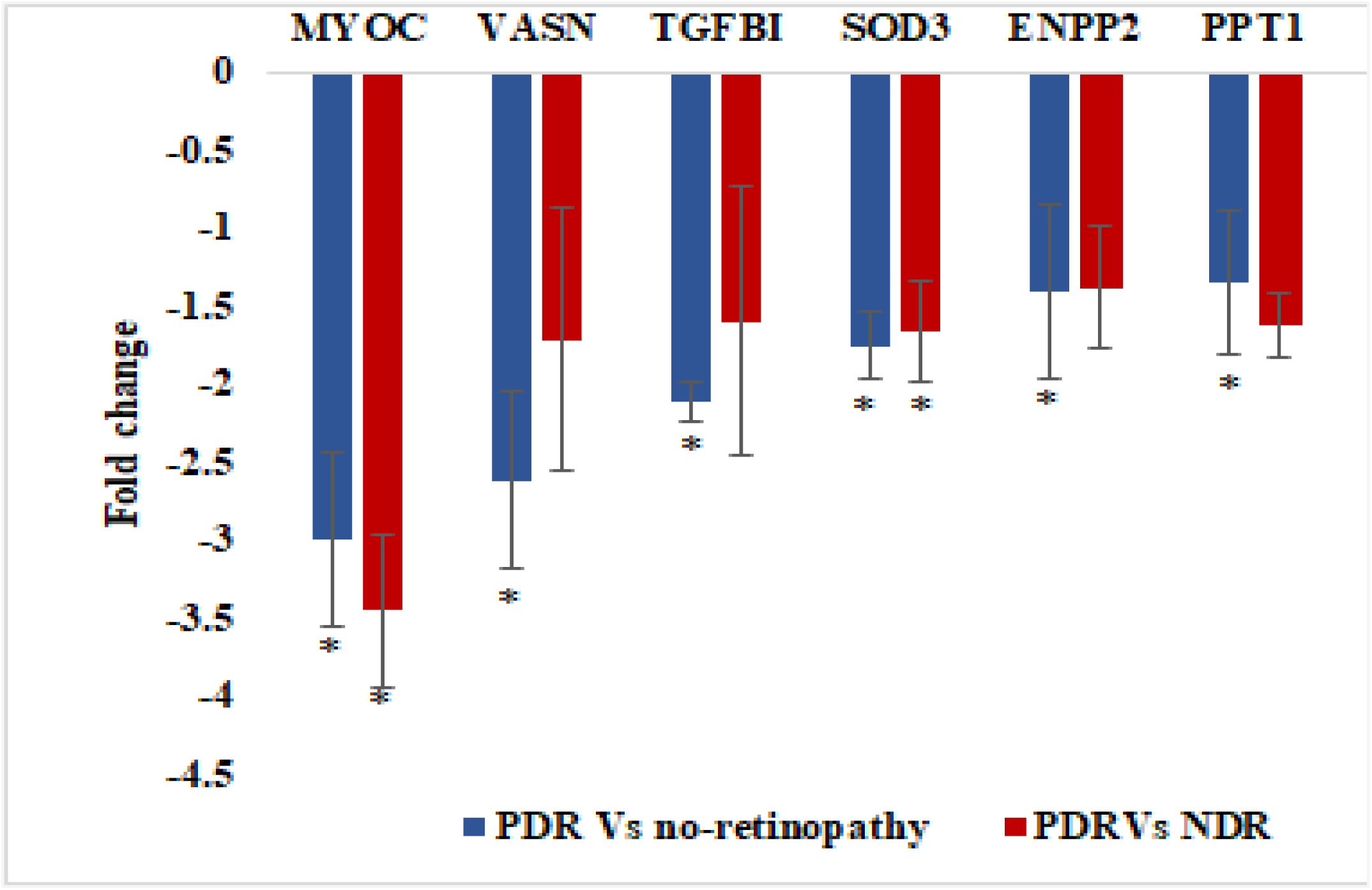
Significantly down-regulated proteins in PDRD vs no retinopathy groups, \**p*<0.05

### Pathway analysis using Ingenuity pathway analysis (IPA) software

The pathway analysis using IPA was performed amongst three categories (DM vs control; PDR vs control and PDR vs DM). The top significant pathways found associated in DM were glycolysis, gluconeogenesis, phagosome maturation pathway, LXR/RXR activation (involved in lipid metabolism, inflammation, cholesterol metabolism), oxidative stress and the hypoxia signaling pathway. 207 molecules were found associated with cellular movement, adhesion, cell death and survival. Alongwith the above pathways, in PDR vs control; coagulation system pathway, natural killer cell signaling, production of ROS in macrophages were also found significantly associated. This signifies increased activation of coagulant factors in PDR blood which is a hallmark of this disease and therefore validates our data/findings. Protein synthesis and degradation and cell-cell signaling were additional molecular functions identified in PDR vs control alongwith the above identified in DM vs control. Angiogenesis, apoptosis, cell death of epithelial cells, binding of myeloid cells, interaction of leucocytes and cell proliferation were the prominent networks identified. TGFB1 was the upstream regulator consistently found associated (inhibition) across all the three data sets. The coagulation and prothrombin activation pathway showed an increasing trend for z-score while we move from control to DM and DM to PDR. While the CLEAR signaling pathway followed a decreasing trend towards PDR. The LXR/RXR activation pathway was most prominent in the PDR vs DM category. The glycolysis and gluconeogenesis were highly significant towards DM and PDR as compared to controls but no effect for the same was observed in PDR vs DM. Therefore, they could be a result of diabetes and not necessarily DR.

### Pathway analysis using R software

Autophagy-associated pathways, inflammation, matrix metalloprotease activity and mitophagy was significantly enriched in our PDR dataset as compared to diabetic individuals (Figure 4).

**Figure 4:**
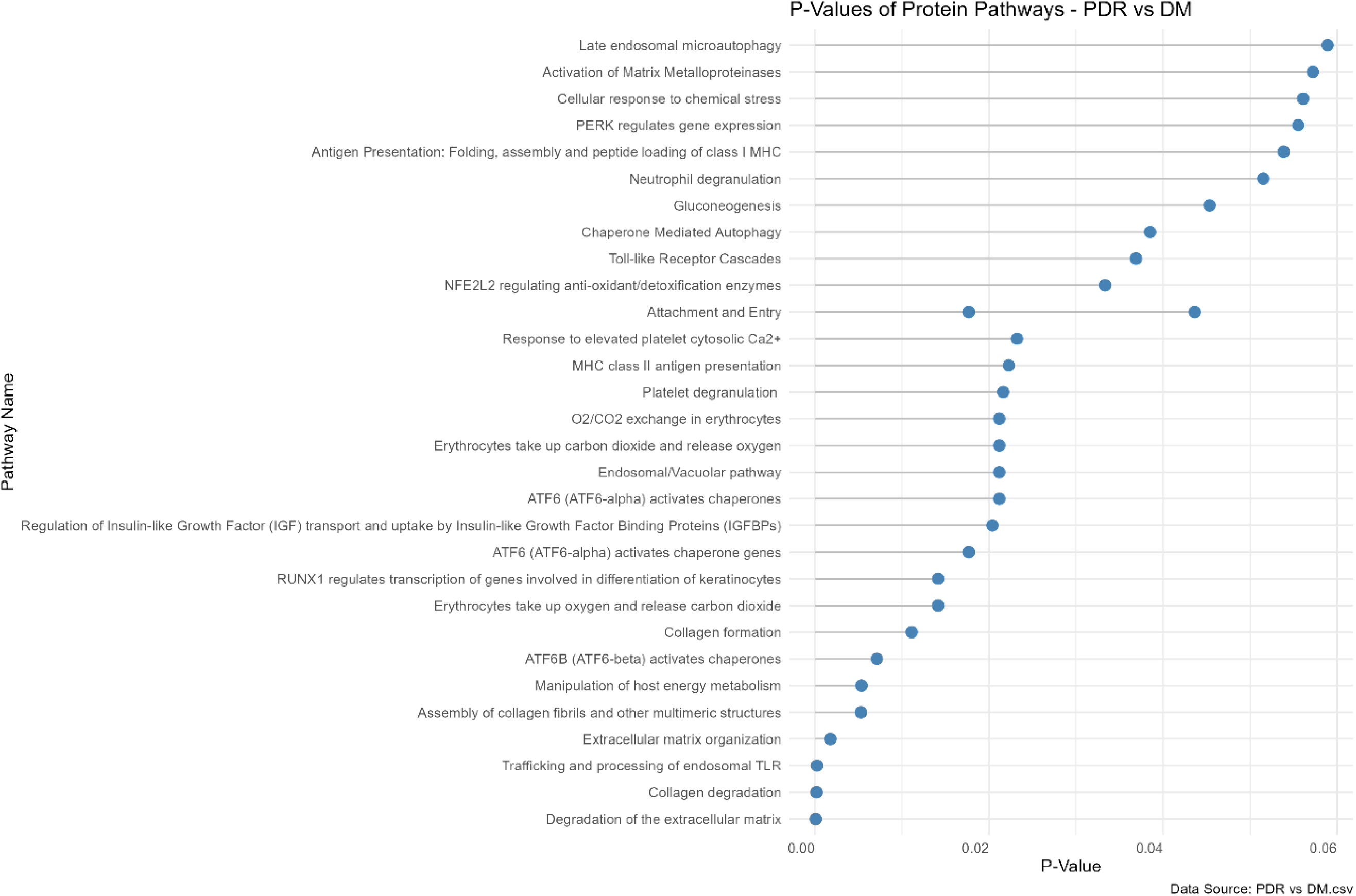
“R” software analysis identifies the given pathways to be significantly enriched in PDR as compared to diabetic individuals.

### Validation of crucial pathways using ELISA

Demographic details of the subjects recruited are provided in the table 4.

**Table 4:** Demographic details of the patient samples used for ELISA.

| Category | No. of samples | Age group (years) | Inclusion criteria | Exclusion criteria |
| --- | --- | --- | --- | --- |
| Control | 25 | 50-65 (mean age 58±7.5) | Cataract, macular hole | Ocular infections, diabetes, thyroid |
| PDR | 25 |  | DM>10 years with PDR changes | Ocular infections |

The validation of the above identified pathways was done by assessing the expression of their pathway proteins using ELISA. The assessed markers were found to be significantly upregulated in the PDR vitreous samples. MMP-9 (inflammatory marker and an regulator of cell death) was almost undetected in the control samples, however it was abundantly present in the PDR vitreous samples (1500pg/mL). IL-8, (an inflammatory marker, neutrophil chemotactic factor widely expressed by macrophages) its mean abundance in controls and PDR vitreous was 150 pg/mL and 450 pg/mL respectively. VEGF, angiogenic marker was also negligibly present in the control vitreous, however had higher expression in the PDR vitreous (1800 pg/mL). Its receptor (VEGFR1) was also significantly increased in the PDR vitreous (920 pg/mL in controls and 1250 pg/mL in PDR). NF-Kb signaling regulates expression of Lipocalin-2 which is a regulator of autophagy, lipid metabolism, neuroinflammation and neurotoxicity was abundantly exressed in the PDR vitreous (1890 pg/mL in controls and 2860 pg/ml in PDR). Mean concentration of thrombospondin;THBS2 (a marker for apoptosis, cell migration, angiogenesis and cytoskeletal regulation) in controls and PDR was 20pg/mL and 4560 pg/mL respectively. The growth factor PDGF-AA (involved in lipid synthesis, cell regulation and growth) was significantly increased in PDR vitreous; 250 pg/mL in controls and 720 pg/mL in PDR vitreous. Expression of endothelin-1 (a vasoconstrictor produced by endothelial cells) was also assessed, though not significant it showed a decreaseing trend in PDR vitreous. LRG1 expression was also assessed to validate the Mass-spec findings in n=10 samples each from DM, NPDR, PDR as compared to controls. It was found to be elevated in both DM as well as PDR vitreous (Figure 5).

**Figure 5:**
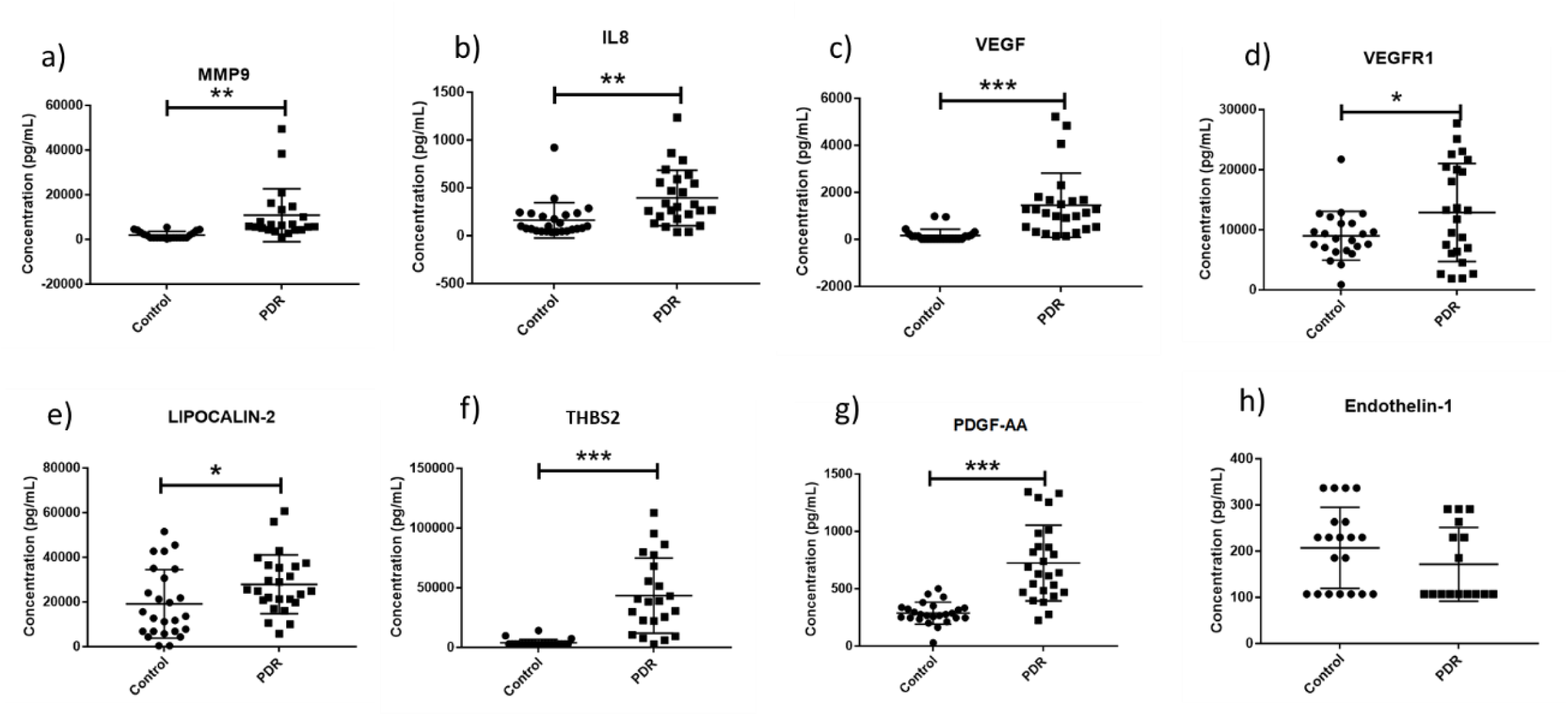
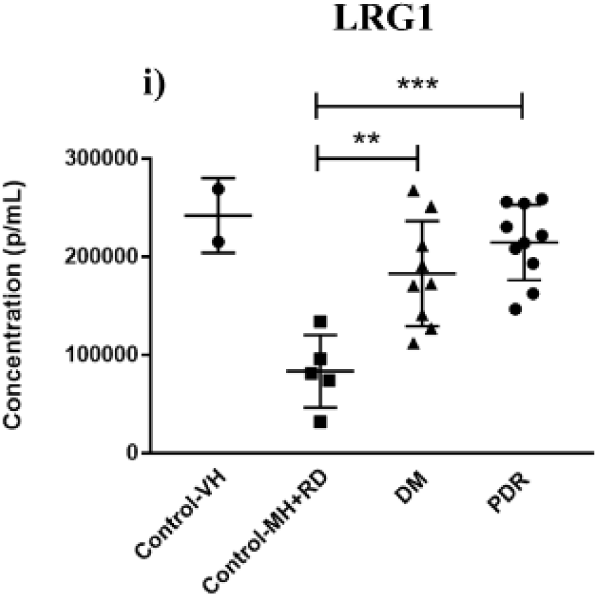
Graphs showing change in protein expression of different proteins assessed using ELISA (n=25; *=p-value <0.05, **=p-value<0.01, ***=p-value<0.001).

### Quantitation of COLIIa and MYOC via western blotting

ColIIa and MYOC expression was validated in the vitreous samples of PDR and controls (n=10 each category) using western blotting. ColIIa was found to be upregulated in the PDR vitreous samples, while MYOC was downregulated as shown in figure 6. Upregulation of ColIIa in vitreous could hint either for extracellular matrix reorganisation or its degeneration leading to secretion of ColIIa in the vitreous cavity. MYOC downregulation signifies inhibition of autophagy in the PDR patients.

**Figure 6:**
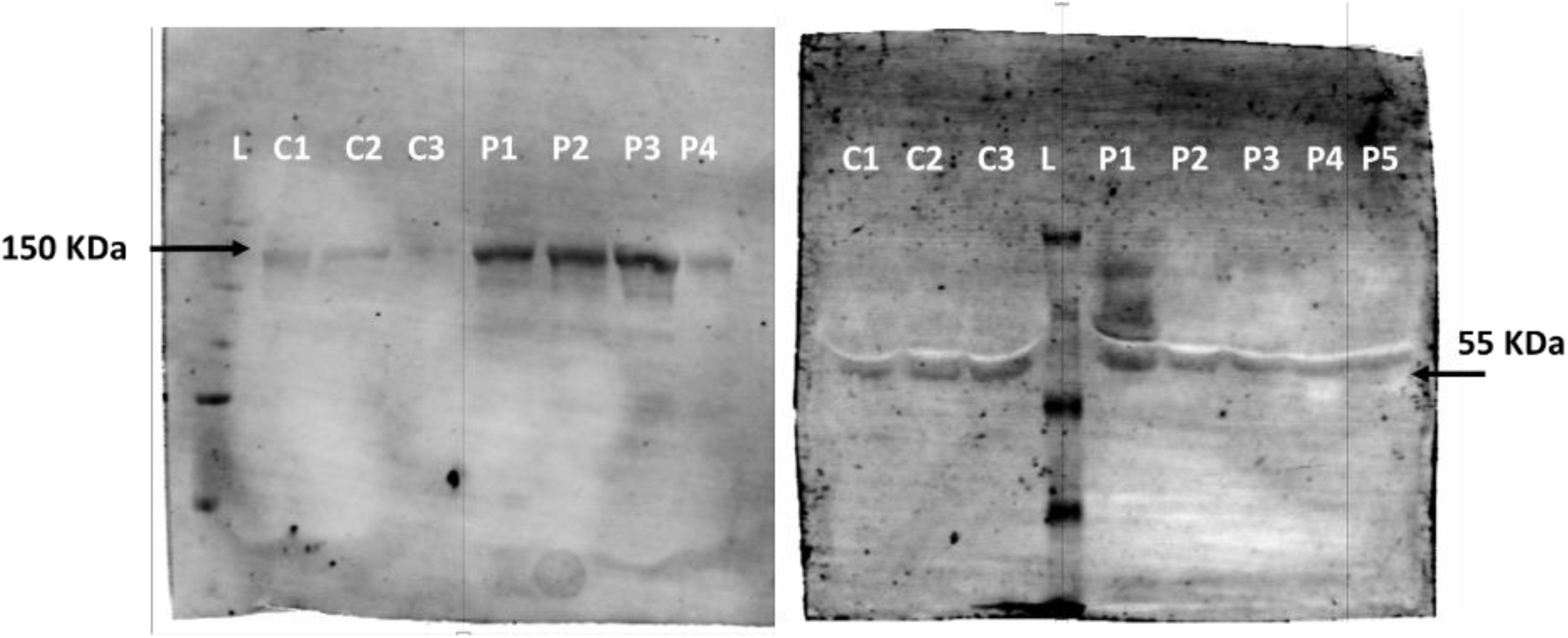
Representative western blots for Col2IIa (left) and MYOC (right) expression in the vitreous samples (n=10 each category). C; Control, PDR; Proliferative DR; L; Ladder.

### COLIIa deposition in the retinal tissues

In order to check collagen deposition surrounding the retinal blood vessels, formalin fixed retinal tissue sections of diabetic and control were subjected to trichome staining. Upon imaging, collagen deposition was observed localised to blood vessels in diabetic retina with higher intensity as compared to the control retina (Figure 7).

**Figure 7:**
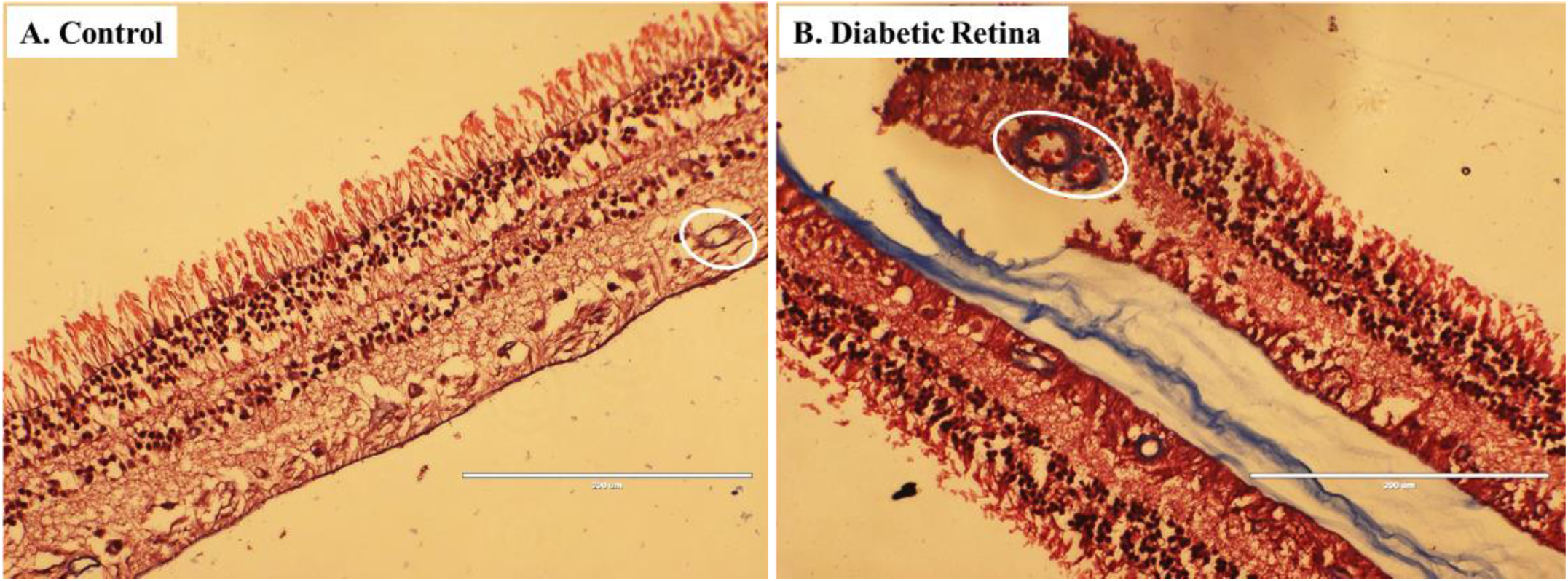
The representative image showing increased collagen deposition (marked with white circle) surrounding the blood vessels of (B) diabetic retina when compared with (A) control retina (n=3; images taken at 20x using EVOS fluorescent microscope).

### TREM2 localisation in the retinal tissues

TREM2 was a novel protein identified in the vitreous of PDR samples. IHC was performed to check for its localisatioin in the PDR and NPDR retinas (n=1 each). The TREM2 protein was found to be expressed in the PDR retinas especially near the blood vessel walls. It was co-localising with the microglial marker F4/80 (yellow region seen in the Figure 8c upper panel). However, the GFAP marker was disntinctly expressed (Figure 8c lower panel).

**Figure 8:**
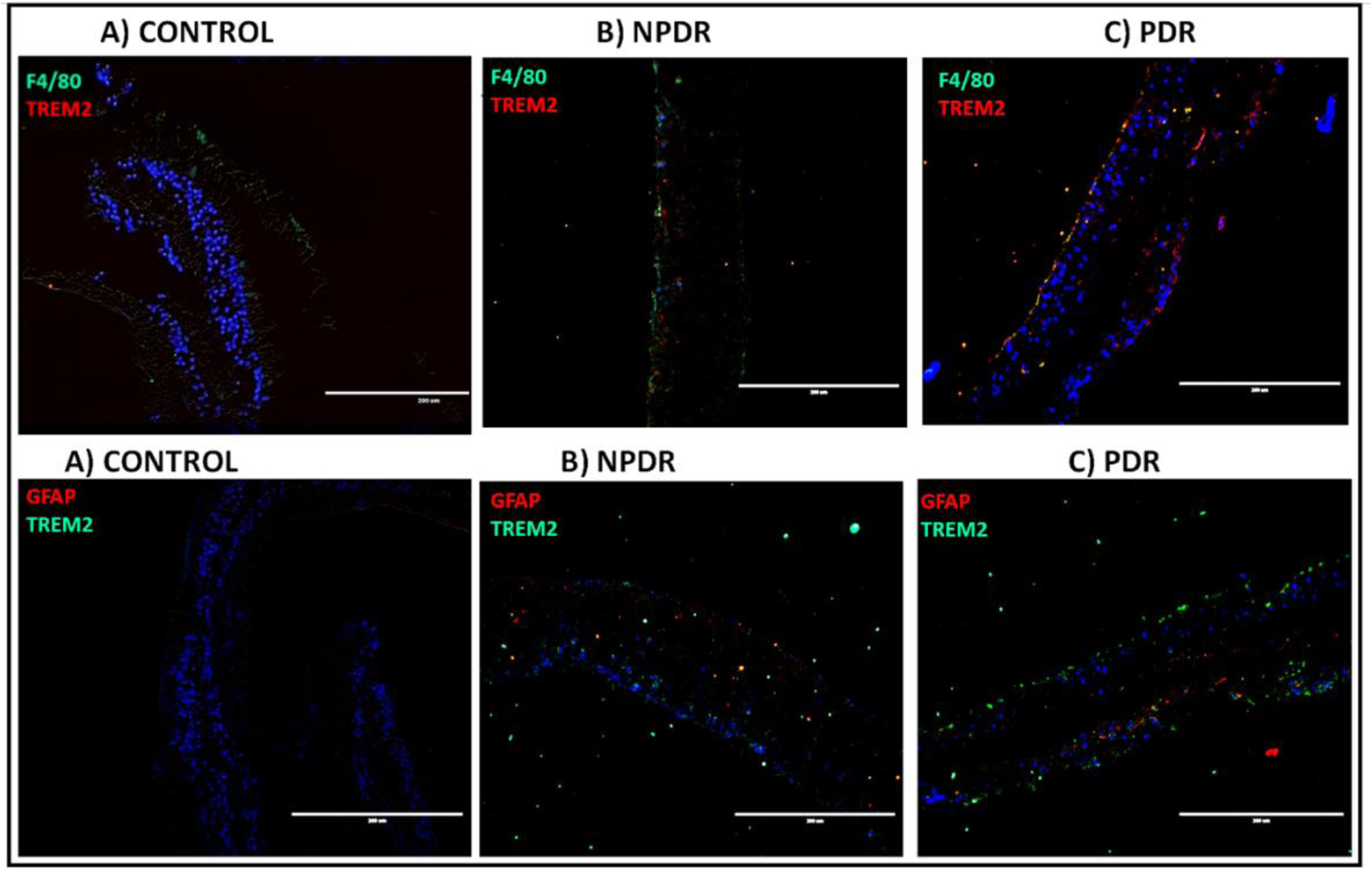
Representative images for IHC in retinal tissues;n=1: a) Control b) NPDR c) PDR for TREM2/ F4/80 and TREM2/ GFAP expression (images taken at 20x using EVOS fluorescent microscope).

### Increased oxidative stress in the PDR environment

SOD3, an antioxidant enzyme was found to be downregulated in the vitreous samples, this signifies increased oxidative stress. Therefore, to confirm that, DCFDA assay was performed to asses for the accumulation of ROS species in the vitreous samples. Upon analysis, DCFDA absorbance was found to be relatively higher in case of the PDR vitreous samples as compared to controls (Figure 9).

**Figure 9:**
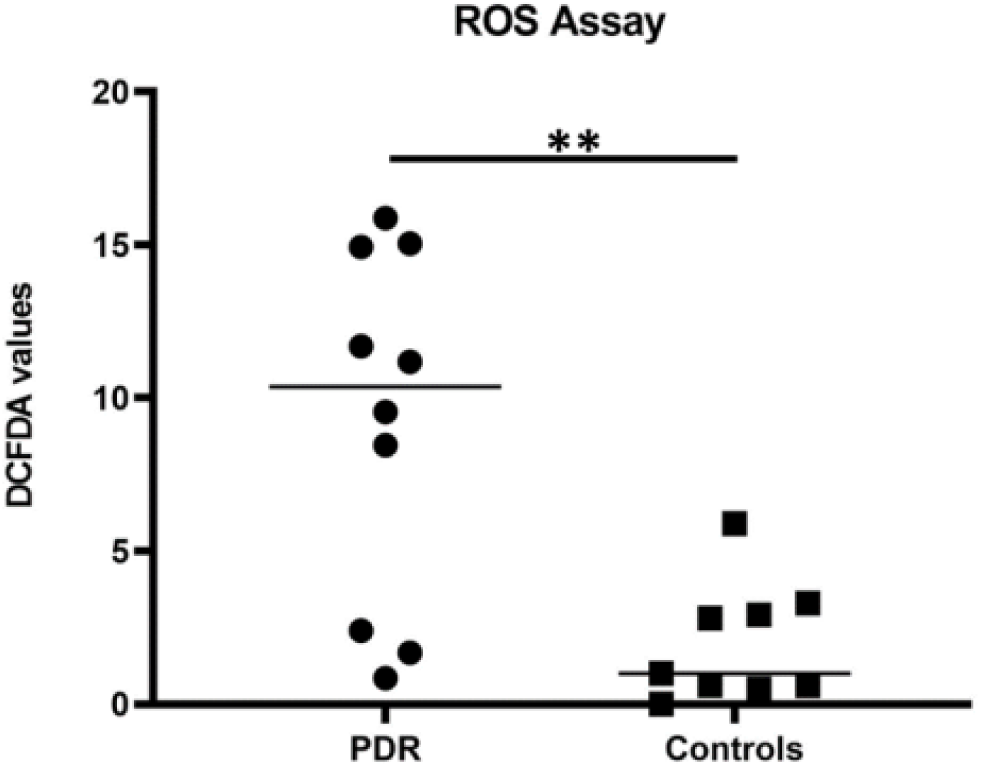
Scatter plot showing DCFDA absorbance in the vitreous samples ** = p-value<0.01.

Differential gene expression of microglial receptors in DR: *TREM2* was observed to be upregulated across all categories and was highly significant in case of PDR vs controls (fc = 3.54; p<0.01). *LDLR* and *APOE* a ligand for TREM2 and another receptor for microglia (APOE is a receptor for astrocytes too) on the other hand was downregulated in DM and NPDR (fc= -1.8, -1.9 and -2.4, -4.3 respectively) but upregulated in case of PDR (fc=1.7, 24.5 respectively) (figure 10).

**Figure 10:**
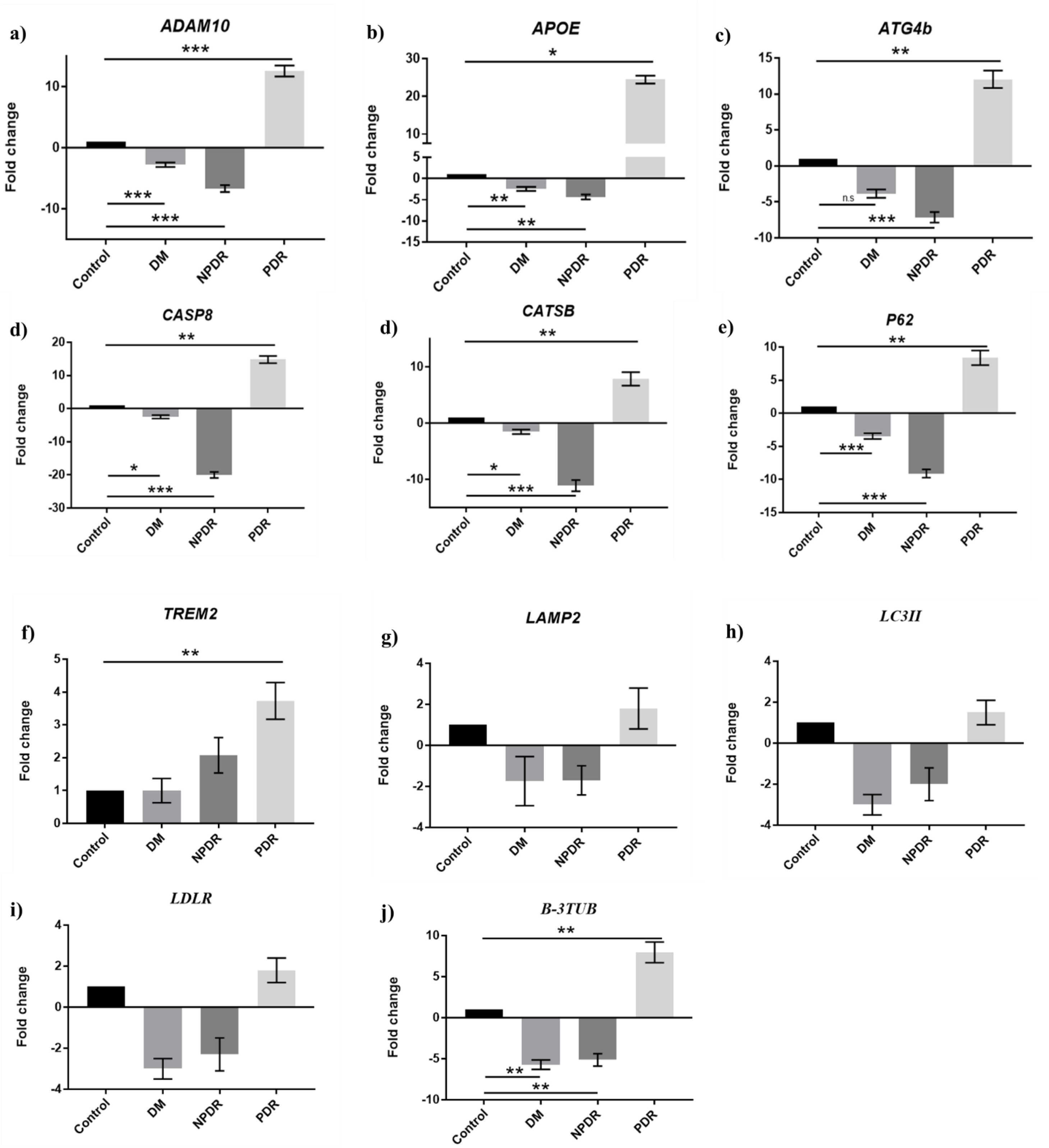
qPCR gene expression analysis for autophagy-associated genes (n= 50 for *TREM2, LDLR, LAMP2* and *LC3II*; n=24 for other genes) in PBMCs isolated from control, DM: Diabetes mellitus, NPDR: Non-proliferative DR and PDR: Proliferative DR patient blood samples. (*=p<0.05, **=p<0.01, ***=p<0.001).

Differential gene expression of autophagy-associated genes in DR: Interestingly, all the assessed genes for autophagy (*ATG4b*; gene essential for autophagosome maturation, *CATSB;* play role in enzymatic degradation of cellular waste, *P62*; delivery of cargo to autophagosome and lysosome, *LAMP2*; required for lysosomal membrane stabilisation and *LC3II;* a protein associated with autophagosome membranes and important for its lipidation during lysosomal fusion) were significantly downregulated in DM and NPDR patient samples. However, the same showed higher upregulation in case of PDR (Figure 10).

Differential gene expression of cellular protease, apoptotic gene and neuronal marker in DR: *ADAM10*, a disintegrin and metalloprotease for microglial receptor TREM2 protein was significantly (p<0.001) downregulated in DM and NPDR (fc = -2, -6 respectively) as compared to controls. But was significantly (p<0.001) upregulated (fc=12) in PDR samples. *CASP8;* a marker for apoptosis and *β-3TUB;* a marker for neuronal biogenesis was also following the same trend (Figure 10).

Differential gene expression of autophagy-associated genes in ocular tissue samples of DR: The expression of all the genes assessed above was validated in ocular tissue samples. mRNA expression of genes was compared among retina of individuals with history of diabetes and age-matched controls. However, to assess the similar phenomenon in DR patients (as availability of retina from DR patients is rare), gene expression was assessed in ERMs which are membranes formed at vitreo-retinal junction in PDR cases and was compared to ERMs collected from age-matched cataract controls. All the assessed genes were found to be downregulated in both the cases (Figure 14). LAMP2 and APOE were significantly (p<0.05) downregulated (fc= -5.2, -10.1 respectively) in the ERMs of PDR compared to controls (Figure 11).

**Figure 11:**
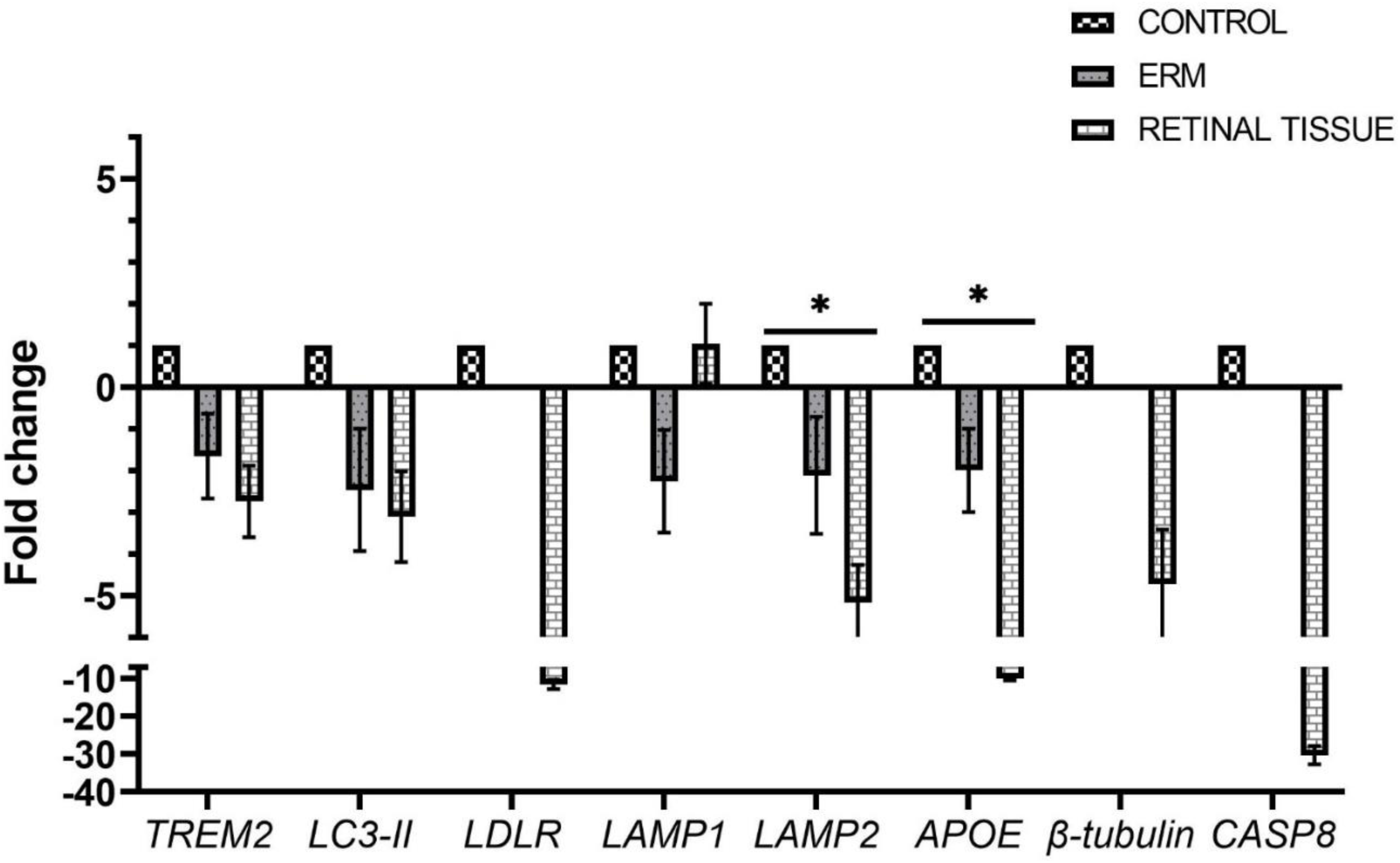
qPCR gene expression analysis for autophagy-associated genes (n= 10; both in retinal tissues and ERMs) isolated from control and DR tissues (*=p<0.05).

Gene expression analysis in primary mixed retinal cell culture exposed to hyperglycaemia: Next, we checked for the expression of these genes in primary mixed retinal cell culture exposed to hyperglycaemia (20Mm D-Glucose). Before performing qPCR, cells were characterised with cell specific markers and cell viability was assessed by alamar blue assay (Figure 12a,12b and figure 12c respectively).

**Figure 12:**
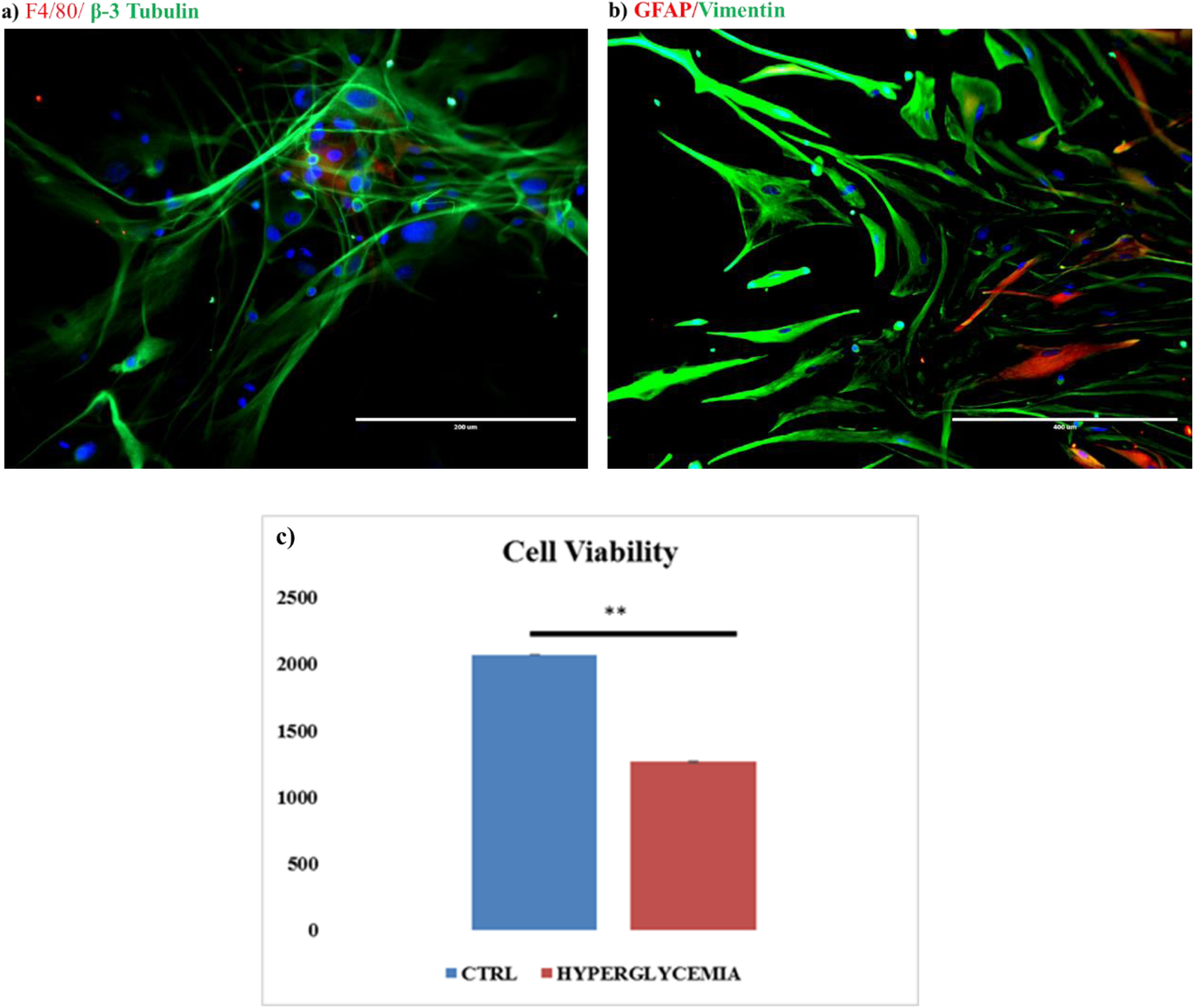
a) F4/80, microglial marker shown in red and β-3 Tubulin a neuronal marker shown in green colour. b) GFAP, a marker for astrocytes and Vimentin, a muller glia marker marked in red and green respectively. “images taken at 20x using EVOS fluorescent microscope”. c) Cell viability assay using alamar blue dye after exposure of PRMCs to 20mM D-glucose for 24 hours (P**<0.01). The y-axis shows number of cells in each category.

QPCR in PRMCs exposed to 20mM D-glucose: The expression of the above genes was also quantified in the PRMCs exposed to hyperglycaemia. After 24 hours of D-glucose treatment, RNA was isolated from cells and reverse transcribed to cDNA and qPCR was performed. The exposure to chronic hyperglycaemia seems to elevate the expression of autophagy-associated genes (Figure 13). The expression of β-3 Tubulin was significantly (p<0.05) upregulated (fc=37.5) indicating increased biogenesis of neurons as a response to stress. Therefore, the PRMCs could serve as a good *in-vitro* model to perform further autophagy associated assays and experiments.

**Figure 13:**
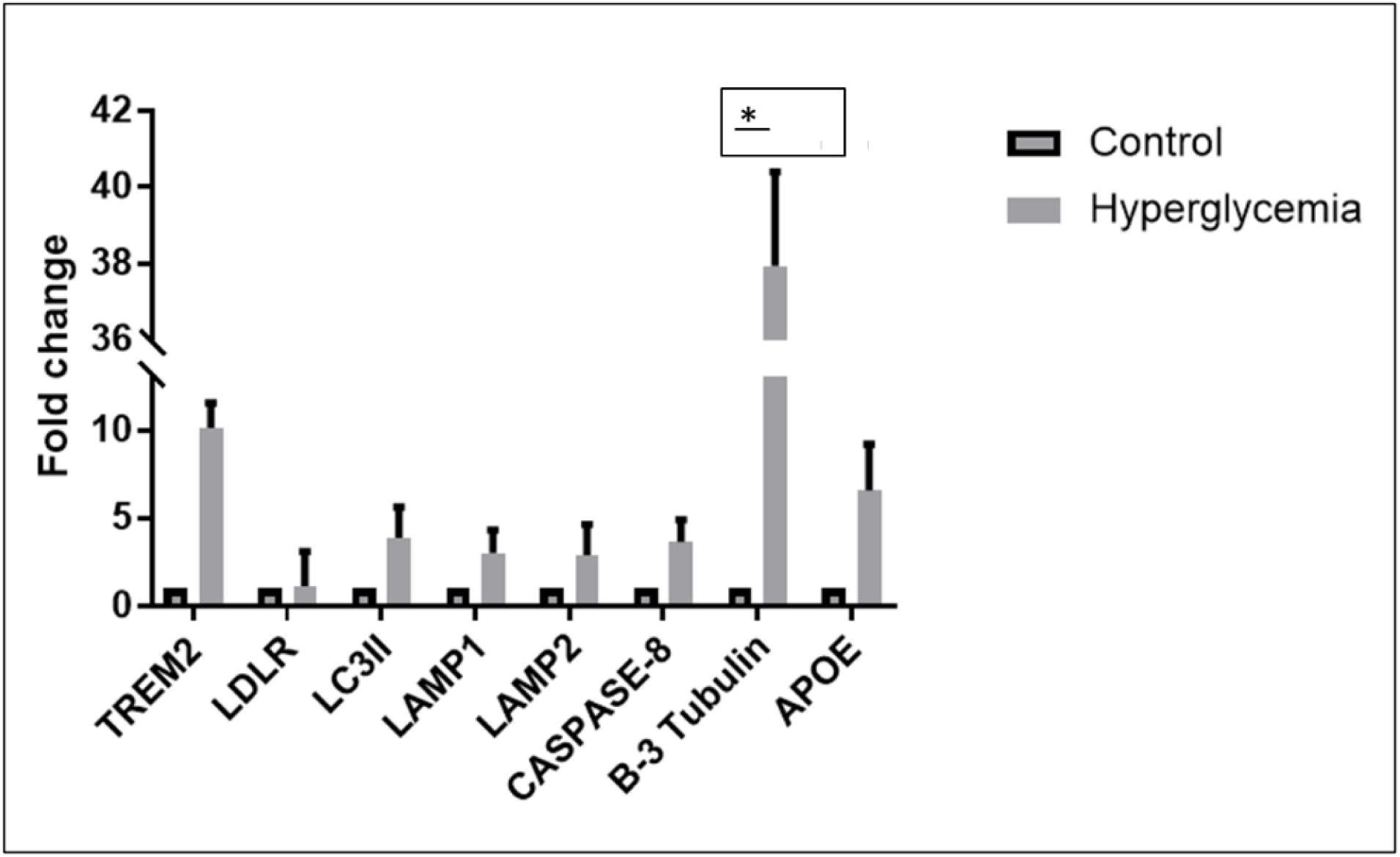
qPCR gene expression analysis for autophagy-associated genes in PRMCs (n=2) *=p<0.05.

## Discussion

Knowledge of protein alterations is required to understand the underlying mechanisms contributing to disease progression in complicated metabolic diseases like DR. Several studies have tried to investigate DR proteome and the related pathways in disease pathology as indicated in the table 5. Vitreous fluid remains the most suitable sample choice for majority of the studies on DR as it is the only accessible ocular fluid, which is most proximally linked to the retina to reflect the underlying proteome changes. However, only a few have compared the alterations of vitreous proteome in DR to those in diabetes without retinopathy (Gao, Chen, Timothy et al. 2008). The comparison of DR proteome with diabetic proteome alone is crucial to identify clearly the retinopathy specific changes. Herein, we attempted to compare the vitreous proteomes of PDR, NDR and NDM to identify key mechansims involved in DR pathogenesis.

**Table 5:**
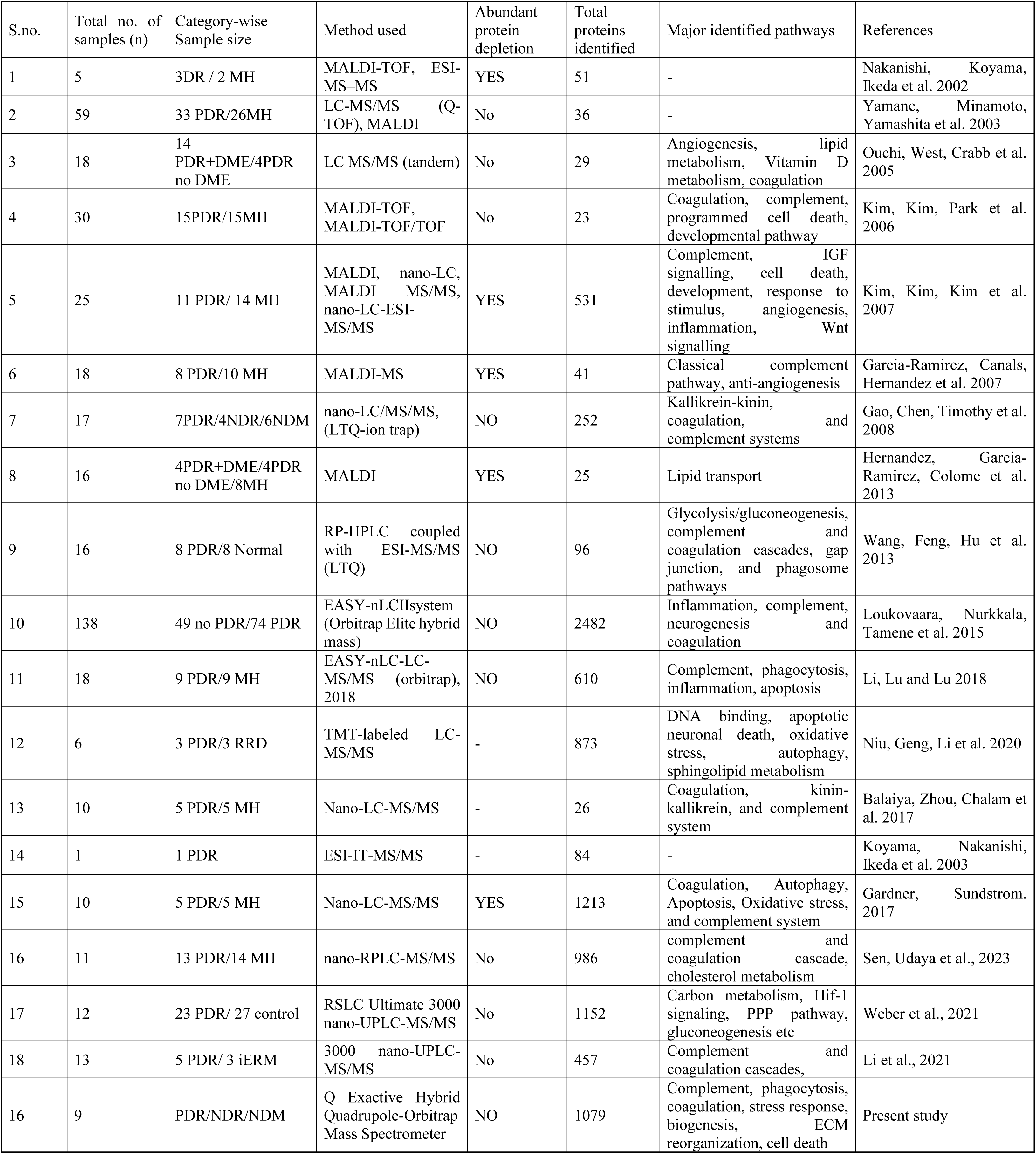
Comparison of vitreous proteome profiling studies in DR by employing different approaches.

| S.no. | Total no. of samples (n) | Category-wise Sample size | Method used | Abundant protein depletion | Total proteins identified | Major identified pathways | References |
| --- | --- | --- | --- | --- | --- | --- | --- |
| 1 | 5 | 3DR / 2 MH | MALDI-TOF, ESI-MS-MS | YES | 51 | - | Nakanishi, Koyama, Ikeda et al. 2002 |
| 2 | 59 | 33 PDR/26MH | LC-MS/MS (Q-TOF), MALDI | No | 36 | - | Yamane, Minamoto, Yamashita et al. 2003 |
| 3 | 18 | 14 PDR+DME/4PDR no DME | LC MS/MS (tandem) | No | 29 | Angiogenesis, lipid metabolism, Vitamin D metabolism, coagulation | Ouchi, West, Crabb et al. 2005 |
| 4 | 30 | 15PDR/15MH | MALDI-TOF, MALDI-TOF/TOF | No | 23 | Coagulation, complement, programmed cell death, developmental pathway | Kim, Kim, Park et al. 2006 |
| 5 | 25 | 11 PDR/ 14 MH | MALDI, nano-LC, MALDI MS/MS, nano-LC-ESI-MS/MS | YES | 531 | Complement, IGF signalling, cell death, development, response to stimulus, angiogenesis, inflammation, Wnt signalling | Kim, Kim, Kim et al. 2007 |
| 6 | 18 | 8 PDR/10 MH | MALDI-MS | YES | 41 | Classical complement pathway, anti-angiogenesis | Garcia-Ramirez, Canals, Hernandez et al. 2007 |
| 7 | 17 | 7PDR/4NDR/6NDM | nano-LC/MS/MS, (LTQ-ion trap) | NO | 252 | Kallikrein-kinin, coagulation, and complement systems | Gao, Chen, Timothy et al. 2008 |
| 8 | 16 | 4PDR+DME/4PDR no DME/8MH | MALDI | YES | 25 | Lipid transport | Hernandez, Garcia-Ramirez, Colome et al. 2013 |
| 9 | 16 | 8 PDR/8 Normal | RP-HPLC coupled with ESI-MS/MS (LTQ) | NO | 96 | Glycolysis/gluconeogenesis, complement and coagulation cascades, gap junction, and phagosome pathways | Wang, Feng, Hu et al. 2013 |
| 10 | 138 | 49 no PDR/74 PDR | EASY-nLCIIsystem (Orbitrap Elite hybrid mass) | NO | 2482 | Inflammation, complement, neurogenesis and coagulation | Loukovaara, Nurkkala, Tamene et al. 2015 |
| 11 | 18 | 9 PDR/9 MH | EASY-nLC-LC-MS/MS (orbitrap), 2018 | NO | 610 | Complement, phagocytosis, inflammation, apoptosis | Li, Lu and Lu 2018 |
| 12 | 6 | 3 PDR/3 RRD | TMT-labeled LC-MS/MS | - | 873 | DNA binding, apoptotic neuronal death, oxidative stress, autophagy, sphingolipid metabolism | Niu, Geng, Li et al. 2020 |
| 13 | 10 | 5 PDR/5 MH | Nano-LC-MS/MS | - | 26 | Coagulation, kinin-kallikrein, and complement system | Balaiya, Zhou, Chalam et al. 2017 |
| 14 | 1 | 1 PDR | ESI-IT-MS/MS | - | 84 | - | Koyama, Nakanishi, Ikeda et al. 2003 |
| 15 | 10 | 5 PDR/5 MH | Nano-LC-MS/MS | YES | 1213 | Coagulation, Autophagy, Apoptosis, Oxidative stress, and complement system | Gardner, Sundstrom. 2017 |
| 16 | 11 | 13 PDR/14 MH | nano-RPLC-MS/MS | No | 986 | complement and coagulation cascade, cholesterol metabolism | Sen, Udaya et al., 2023 |
| 17 | 12 | 23 PDR/ 27 control | RSLC Ultimate 3000 nano-UPLC-MS/MS | No | 1152 | Carbon metabolism, Hif-1 signaling, PPP pathway, gluconeogenesis etc | Weber et al., 2021 |
| 18 | 13 | 5 PDR/ 3 iERM | 3000 nano-UPLC-MS/MS | No | 457 | Complement and coagulation cascades, | Li et al., 2021 |
| 16 | 9 | PDR/NDR/NDM | Q Exactive Hybrid Quadrupole-Orbitrap Mass Spectrometer | NO | 1079 | Complement, phagocytosis, coagulation, stress response, biogenesis, ECM reorganization, cell death | Present study |

A comparison of the identified proteins in the present study with human vitreous proteome database generated as a part of human eye proteome project published in the year 2013 and 2018 (Semba et al., 2013; Ahmad et al., 2018) and those not listed in the vitreous proteome from the eye proteome database (Li et al., 2018; Santos et al., 2018; Zou et al., 2018a; Zou et al., 2018b showed 938 of 1079 proteins were already reported, which confirmed the validity of the data generated. Following stringent criteria including the identification of proteins in a minimum of three vitreous samples with minimum sequence coverage of 2%, 27 novel vitreous proteins were identified in this study. Among these, 16 proteins were reported for the first time (Table 03), whereas the remaining 11 were identified in the eye proteome database (Table 2) (Semba et al., 2013; Ahmad et al., 2018). Eight of these proteins were found to be immunoglobulins and two HLA proteins such as HLA class I histocompatibility antigen-Cw-12 alpha chain (HLA-C) and HLA class I histocompatibility antigen-Cw-17 alpha chain (HLA-C). Two novel vitreous proteins identified were found to be associated with nervous system including TREM2 and VAMP1. TREM2 is a myeloid cell receptor that is expressed exclusively in microglial cells. Activation of this receptor in microglial cells promote microglial activation and phagocytosis and thus plays a critical role in neurodegenerative diseases (Mecca et al., 2018). Even though the role of TREM2 was not explored in DR or diabetes associated complications, its role in Alzheimer’s disease in activating microglia and promoting inflammation has already been reported (Geerlings et al., 2017). VAMP1 is a nervous system-specific protein, which is involved in synaptic functions mainly in the neurotransmitter transport and release (Trimble et al., 1988).

As demonstrated in fig. 2a, most of the elevated proteins in the DM group Agrin (AGRN), COL1A2 (collagen type I alpha 2 chain), and TIMP2 were involved in ECM remodeling, rearrangement, and destruction. Cell survival and neuronal outgrowth are affected by APLP1 (amyloid like protein 1) and APLP2 (amyloid like protein 2). (Sakai and Hohjoh 2006). Insulin-like growth factor-binding protein (IGFBP) is a protein that causes changes in the retina and vitreous, and its expression was increased in the diabetes group in the present study. IGFBP6 alongwith VCAN, a proteoglycan involved in cell motility, proliferation, and differentiation as well as inflammation, were also shown to be elevated in diabetic subjects (Wight and Merrilees 2004). Simultaneously, the expressions of 7 proteins were downregulated in the diabetic group. These included the proteins involved in detoxification mechanism and metabolic proteins (Fig 2b). Some studies had shown PRDX1 upregulation as an inflammatory stimulus in various body complications as it enhanced the inflammatory signals like NFkB (Ishii, Warabi and Yanagawa 2012), while others reported that PRDX1 protected cells from oxidative stress induced damage (Kim, Park, Choi et al. 2015). The present study identified a significant downregulation of this protein in the DM, suggesting that the downregulation of PRDX1 and GSTP1 enhanced the oxidative stress induced damage in diabetic complications. Vasorin (VASN), a TGF-β binding protein was found to be significantly downregulated in diabetes as well as in the retinopathy group (DM vs NDM: -2.66±0.36, p=0.01, PDR vs no retinopathy: -2.61±0.57, p=0.01). It modulates the vascular response to injury by inhibiting the level of TGFβ and thereby prevents fibrosis as seen in vascular injury. Its downregulation seems to aid in the fibroproliferative response to vascular damage (Ikeda et al. 2004).

Further it was important to understand the differential proteomic signatures of PDR VS NDR group. We found 10 proteins which were differentially regulated (4 upregulated and 6 downregulated) among these categories (Fig 3a and 3b). The upregulated proteins were angiogenic proteins (LRG1, TGFβ1) and proteins associated with autoimmune and neurodegenerative diseases such as APCS (Serum amyloid-p-component) and CRP (C-reactive protein) (Wang, Abraham, McKenzie et al. 2013b). FGA (a coagulation protein) was also found upregulated in PDR vitreous. On the other hand, SOD3 (a detoxification enzyme) was found to be downregulated in PDR vitreous. Its downregulation suggests defects in antioxidant mechanism in PDR condition which further deteriorates the ocular environment. Downregulation of fibronectin binding protein, myocilin (MYOC), a protein involved in increased resistance to apoptosis was seen in PDR (Joe, Kwon, Cojocaru et al. 2014). This could explain increased apoptotic cell death in diabetic retinopathy individuals. Lysosomal proteins (ENPP2 and PPT1) were also downregulated in PDR vitreous. Palmitoyl-protein thioesterase 1 (PPT1) is involved in the lysosomal mediated degradation of tagged proteins. The downregulation of these lysosomal enzymes suggests defects in autophagy. Role of autophagy and autophagic proteins has not been defined in DR yet. Accumulation of reactive oxygen species are a known inducer of autophagy and was evident in our PDR subjects (Figure 9). But could be the hallmark of early diabetic retinopathy changes such as cell death seen in individuals.

The GO-annotation of identified proteins is used to determine the associated molecular and biological processes. Immune activation is known to be implicated in DR pathology and the same has been confirmed by this study too. Number of immune system proteins were identified which exhibited a gradual increase from diabetes to PDR, validating earlier research’ findings that immune activation plays a role in PDR (Rajab, Baker, Hunt et al. 2015, Xu and Chen 2017). An increased phagocytic biological process in the PDR group as compared to NDR and NDM suggests that cell death and related mechanisms like autophagy and apoptosis are involved in the pathogenesis of PDR. This becomes interesting as the present study also reports the presence of novel protein TREM2 (involved in microglial autophagy) in the vitreous proteome. Therefore, we have further demonstrated here the role of TREM2 in retinal neurodegeneration. In our data, TREM2 gene was found to be upregulated in the PBMCs of different categories of DR patients (Fig.10F). This is in concordance with the study published in the year 2018 where authors observed increased expression of TREM2 in PBMCs of Alzheimer patients. This was explained by the fact that the expression increases as a counteraction to inhibit neuroinflammation (Casati M *et al*, 2018). Alongwith increase inTREM2 gene expression, there was concurrent decrease in the other genes in the NPDR category whereas it was upregulated in PDR (Fig 10). Decreased expression of TREM2 ligands such as LDLR and APOE in NPDR suggests reduced TREM2 signalling which could lead to reduced microglial phagocytosis and autophagy. This is also evident from the downregulation of autophagy proteins in NPDR suggesting decreased autophagy in the mild form of the disease. Decreased autophagy could lead to increased apoptosis which is evident in DM as well as NPDR retina. ADAM10 is a metalloprotease that cleaves the transmembrane domain of TREM2 to form soluble TREM2 (sTREM2). sTREM2 plays role in ligand binding and microglial activation. This signifies reduced activation of microglia in case of NPDR which can also mean reduced autophagy. Reduced biogenesis of neurons in case of NPDR as evident from reduced expression of β-3 Tubulin. Caspase-8 is pro-apoptotic and inflammatory marker but also protects against a type of cell death known as necroptosis. Its reduced expression in NPDR suggests induction of necroptotic cell death in case of NPDR. Interestingly, all these genes were found upregulated in the PDR category (Fig 10). Now, in case of PDR the vascular abnormalities are dominant over the neurodegenerative changes. There is increased proliferation of vascular endothelial cells in response to ischemia in case of PDR. Autophagy is also involved in the induction of collagen production leading to collagen deposition which was evident in our data (Figure 6). The collagen deposition was also confirmed in the diabetic retinas suggesting a key role of autophagy in early DR development (Figure 7). To conclude, our study findings implicate autophagy besides other well-known mechanistic pathways in DR that could potentially serves as a therapeutic target after careful validation among different cohorts worldwide.

## Limitations

Smaller sample size for mass spec proteomic analysis is a major limitation of this study, however, this was minimized by validation of the data at various levels and by using different techniques. Due to rarity of human retinal tissues from DR tissues, immunohistochemistry was performed only in a single sample and thus warrants further validation.

## Acknowledgement

We acknowledge Ramayamma International Eye Bank for providing us with the donor eyeballs. We thank Hyderabad Eye research Foundation and LV Prasad Eye Institute for providing us with the infrastructural support. We thank patients for the valuable samples, and our pathology team for their help with tissue sectioning work.

